# Interplay between nutrient transporters ensures fertility in the malaria mosquito *Anopheles gambiae*

**DOI:** 10.1101/2023.06.02.543516

**Authors:** Iryna Stryapunina, Maurice Itoe, Queenie Trinh, Charles Vidoudez, Esrah Du, Lydia Mendoza, Oleksandr Hulai, Jamie Kauffman, John Carew, William Robert Shaw, Flaminia Catteruccia

## Abstract

Females from many mosquito species feed on blood to acquire nutrients for egg development. The oogenetic cycle has been characterized in the arboviral vector *Aedes aegypti*, where after a bloodmeal, the lipid transporter lipophorin (Lp) shuttles lipids from the midgut and fat body to the ovaries, and a yolk precursor protein, vitellogenin (Vg), is deposited into the oocyte by receptor-mediated endocytosis. Our understanding of how the roles of these two nutrient transporters are mutually coordinated is however limited in this and other mosquito species. Here, we demonstrate that in the malaria mosquito *Anopheles gambiae*, Lp and Vg are reciprocally regulated in a timely manner to optimize egg development and ensure fertility. Defective lipid transport via *Lp* silencing triggers abortive ovarian follicle development, leading to misregulation of Vg and aberrant yolk granules. Conversely, depletion of Vg causes an upregulation of *Lp* in the fat body in a manner that appears to be at least partially dependent on target of rapamycin (TOR) signaling, resulting in excess lipid accumulation in the developing follicles. Embryos deposited by Vg-depleted mothers are completely infertile, and are arrested early during development, likely due to severely reduced amino acid levels and protein synthesis. Our findings demonstrate that the mutual regulation of these two nutrient transporters is essential to safeguard fertility by ensuring correct nutrient balance in the developing oocyte, and validate Vg and Lp as two potential candidates for mosquito control.

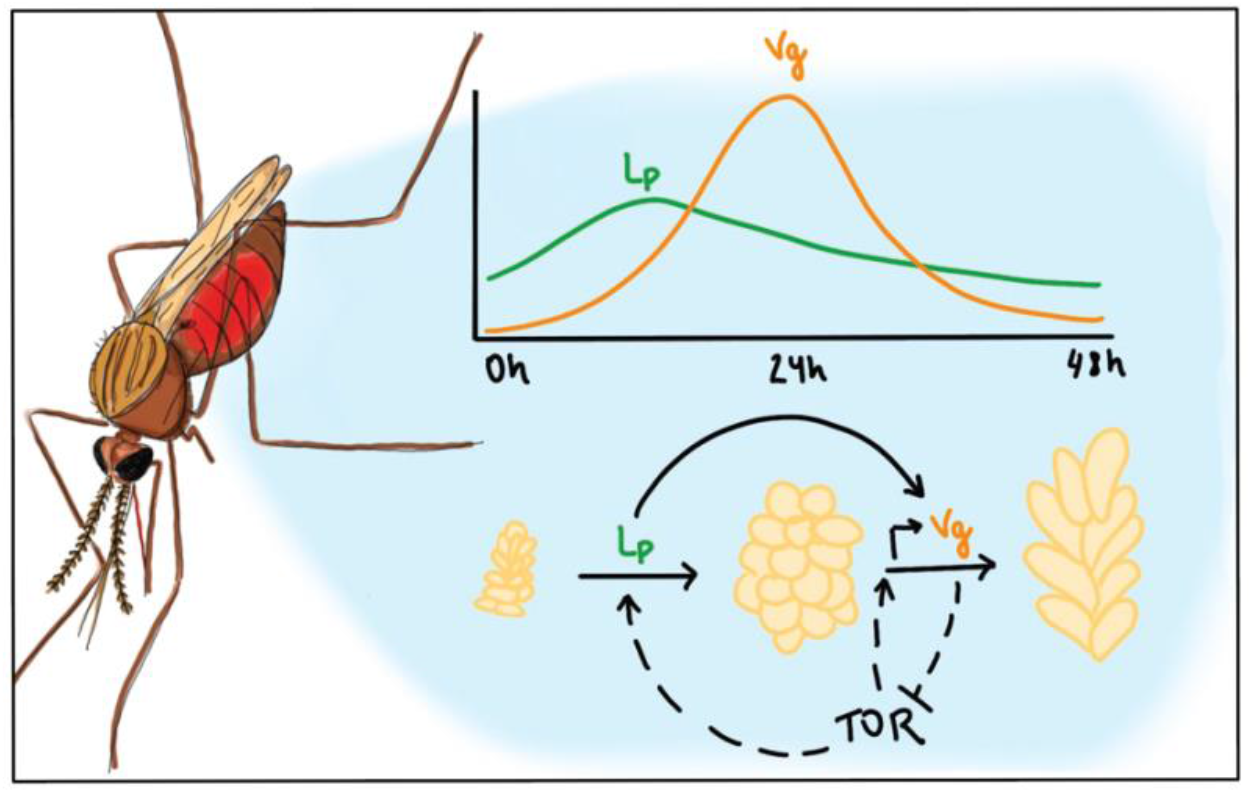

## Introduction

The *Anopheles gambiae* mosquito is one of the most important vectors for the transmission of *Plasmodium falciparum*, a malaria parasite that causes remarkable morbidity and mortality in sub-Saharan Africa and other tropical and subtropical regions (World Malaria Report 2022). Transmission starts when a female mosquito takes a blood meal from a human infected with the sexual stages of *Plasmodium*. At the same time as parasite development begins in the midgut, females use blood nutrients to start their reproductive cycle, which culminates in the development of a full set of eggs in about 2-3 days. The signalling cascades triggered by blood feeding and leading to successful egg development have been largely elucidated in *Aedes aegypti* mosquitoes. In this species, the ovarian ecdysteroidogenic hormone (OEH) and insulin-like peptides (ILPs) are released from the brain upon blood feeding, stimulating the production of the ecdysteroid ecdysone (E, which is synthetized from cholesterol) by the ovarian epithelium (Brown et al., 1998; Brown et al., 2008). After transport to the fat body, E is converted to 20-hydroxyecdysone (20E, the active form of this steroid hormone), which binds to its nuclear receptor to trigger transcriptional cascades leading to the activation and repression of hundreds of genes. Among these genes is *Vitellogenin* (*Vg*), the main egg yolk protein precursor in oviparous species, which peaks at 24 hours post blood meal (PBM) (Attardo, Hansen, and Raikhel 2005). Vg is then released from the fat body into the hemolymph, from where it is taken up by the ovaries by receptor-mediated endocytosis (Roth and Porter 1964; Raikhel and Dhadialla 1992). In the oocytes Vg is crystalized into vitellin, which forms the yolk bodies that the embryo uses as a nutritional source of amino acids (Clements 1992; Li and Zhang 2017; Kunkel and Nordin 1985).

Prior to *Vg* expression, the lipid transporter lipophorin (Lp) shuttles cholesterol and neutral lipids (mostly triglycerides (Ford and Van Heusden 1994)) from the midgut to the ovaries, starting the early phase of egg development and the synthesis of the steroid hormone E (Sun et al. 2000; Atella et al. 2006). It is unclear how *Lp* expression is regulated, although *ex vivo* experiments in *Ae. aegypti* have shown this lipid transporter to be upregulated upon fat body exposure to 20E (Sun et al. 2000). After egg development is completed, if the female is mated, she will oviposit her eggs and return to the pre-blood meal metabolic state. At this point she is ready to begin another gonotrophic cycle, consisting of blood feeding, oogenesis and oviposition.

Besides triggering the synthesis of 20E through cholesterol uptake and E release by the ovaries, blood meal digestion and the subsequent influx of amino acids and ILPs results in the activation of the target of rapamycin (TOR) signalling pathway (Attardo, Hansen, and Raikhel 2005; Hansen et al. 2014). The integration of these nutritional signals leads to a TOR-mediated global regulation of translation and transcription of specific genes that control growth and metabolism (Valvezan and Manning 2019), including *Vg* transcription in mosquitoes (Hansen et al. 2004; reviewed in Attardo, Hansen, and Raikhel 2005; Hansen et al. 2014). TOR regulates translation by directly phosphorylating S6 kinase (S6K) (Hansen et al. 2005), which in turn phosphorylates S6, thus regulating translation of ribosomal proteins and translation elongation factors (reviewed in Attardo, Hansen, and Raikhel 2005; Hansen et al. 2014). S6K also activates the translation of AaGATAa, which promotes *Vg* transcription in *Ae. aegypti* (Park et al. 2006). Due to its central role in the integration of nutritional signals post blood meal, abrogation of TOR signalling results in reduced fecundity in *Ae. aegypti* and *Anopheles stephensi* (Hansen et al. 2004; 2005; Wang et al. 2022; Feng et al. 2021).

Although successful oogenesis in mosquitoes is likely to be tightly dependent on the coordinated function of Lp and Vg, very limited information is available concerning whether and how these nutrient transporters are mutually regulated to ensure egg development and fertility. A study conducted in *An. gambiae* showed that *Lp* knockdown results in reduced expression of *Vg* after feeding on mice infected with the rodent malaria parasite *Plasmodium berghei*, while *Vg* silencing did not affect *Lp* expression (Rono et al. 2010). No further studies have clarified the possible co-regulation of these factors in ensuring accurate nutrient deposition during oogenesis. Additionally, data concerning the mechanisms regulating egg development in *An. gambiae* are sparce despite the key importance of this species for malaria transmission. In these mosquitoes, silencing of *Lp* (whose expression peaks at 12-18h PBM) has been shown to severely hamper egg development (Vlachou et al. 2005; Rono et al. 2010; Werling et al. 2019), revealing a similar role to *Aedes*. Interestingly, *Lp* levels were upregulated upon inhibition of 20E signalling via silencing of the nuclear *EcR* receptor, suggesting that in certain conditions 20E may repress its expression (Werling et al. 2019). The role of Vg in *An. gambiae* reproduction has been studied even more marginally, with a single study reporting that its depletion results in fewer females developing mature oocytes (Rono et al. 2010).

Here we show that the functions of Lp and Vg are tightly linked in *An. gambiae*. Using functional knockdowns of these genes combined with electron microscopy, multi-omics and biochemistry analyses, we show that depletion of these factors results in profoundly deleterious effects on fecundity and fertility. While Lp is required for successful egg development, Vg is essential for fertility as its depletion leads to an early block in embryonic development. We also prove that the functions of these factors are mutually co-regulated, as Lp is needed for the correct incorporation of Vg in the developing oocytes while Vg is required to terminate the Lp-mediated accumulation of lipids in the ovaries. Intriguingly, our data suggest that both induction of *Vg* after blood feeding and its downstream effects on Lp are mediated by TOR signalling, which in turn appears to be repressed by Vg following a blood meal. Our data reveal an intricate reproductive system based on the timely and mutually coordinated function of these nutrient transporters, and identify novel potential targets to interfere with the fertility of these important malaria vectors.

## Results

### *Lp* knockdown significantly impairs oogenesis and affects Vg localization

Previous studies have shown that *Lp* silencing results in severely reduced oogenesis in *An. gambiae* females (Vlachou et al. 2005; Rono et al. 2010; Werling et al. 2019) but without characterizing the phenotype. We confirmed these findings by injecting double stranded RNA targeting *Lp* (dsLp) prior to blood feeding (Supplementary Figure 1A), which dramatically reduced the median number of eggs developed by females compared to controls (injected with dsLacZ) (Figure 1A). This reduction in egg numbers was characterized by a striking decrease in triglycerides in the ovaries at different time points post blood meal (PBM), paralleled by a substantial accumulation of these lipids in the midgut, consistent with a role of Lp in shuttling these lipids from the midgut to the developing eggs (Figure 1B).

**Figure 1.**
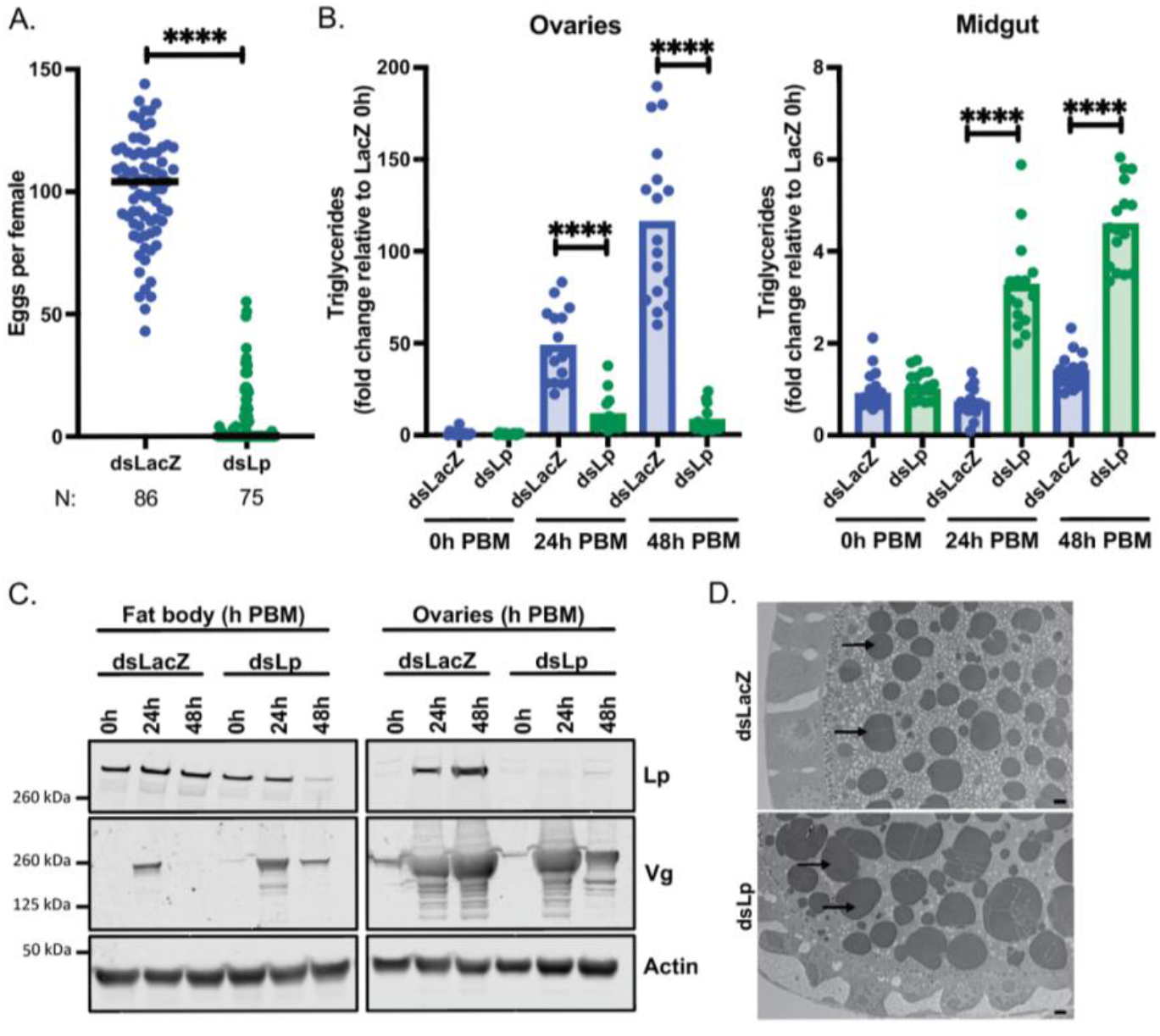
*Lp* knockdown significantly impairs oogenesis and affects Vg localization. (A) Following *Lp* knockdown, females develop a decreased number of eggs; each dot represents eggs per female; N = number of females (Mann-Whitney: **** = p < 0.0001; four biological replicates). (B) Triglycerides accumulate in the midgut of Lp-depleted females, and fail to accumulate in the ovaries; PBM = post blood meal (REML variance component analysis by timepoint: **** = p < 0.0001; three biological replicates). (C) Vg is persistently detectable in the fat body of Lp-depleted females and is decreased in the ovary at 48h PBM (three biological replicates). (D) Transmission electron microscopy showing larger yolk granules (arrows pointing at darkly staining circles) in the ds*Lp* ovary compared to ds*LacZ* at 24h PBM; scale bar = 2µm (one biological replicate).

We next examined how *Lp* knockdown affects levels of *Vg*, the major nutrient transporter incorporated into developing eggs. Interestingly, *Lp*-silenced females had aberrant Vg production and distribution. Not only was *Vg* expressed at lower levels upon Lp depletion (Supplemental Figure 1B), consistent with previous work (Rono et al. 2010), but also this yolk protein showed atypical localization. While in control samples Vg was detected in the fat body at 24h PBM and was fully incorporated into the ovaries by 48h PBM, in *Lp*-silenced females Vg persisted in the fat body at the latter time point and its levels were reduced in the ovaries (Figure 1C; Supplemental Figure 1C). This was consistent with our observation that the ovaries of Lp-depleted females develop normally up to 24h PBM but degenerate by 48h PBM (Supplemental Figure 1D). Interestingly, as observed by electron microscopy, yolk granules in the few eggs that develop after *Lp* silencing appeared larger at 24h PBM than those in control ovaries, and Vg had largely disappeared from western blots by 48h PBM (Figure 1D), suggesting yolk degradation at this latter time point.

Together, these data show that Lp-mediated shuttling of lipids from the midgut into the ovaries is an essential check point for oogenesis, as preventing this process results in misregulated vitellogenesis.

**Supplemental Figure 1.**
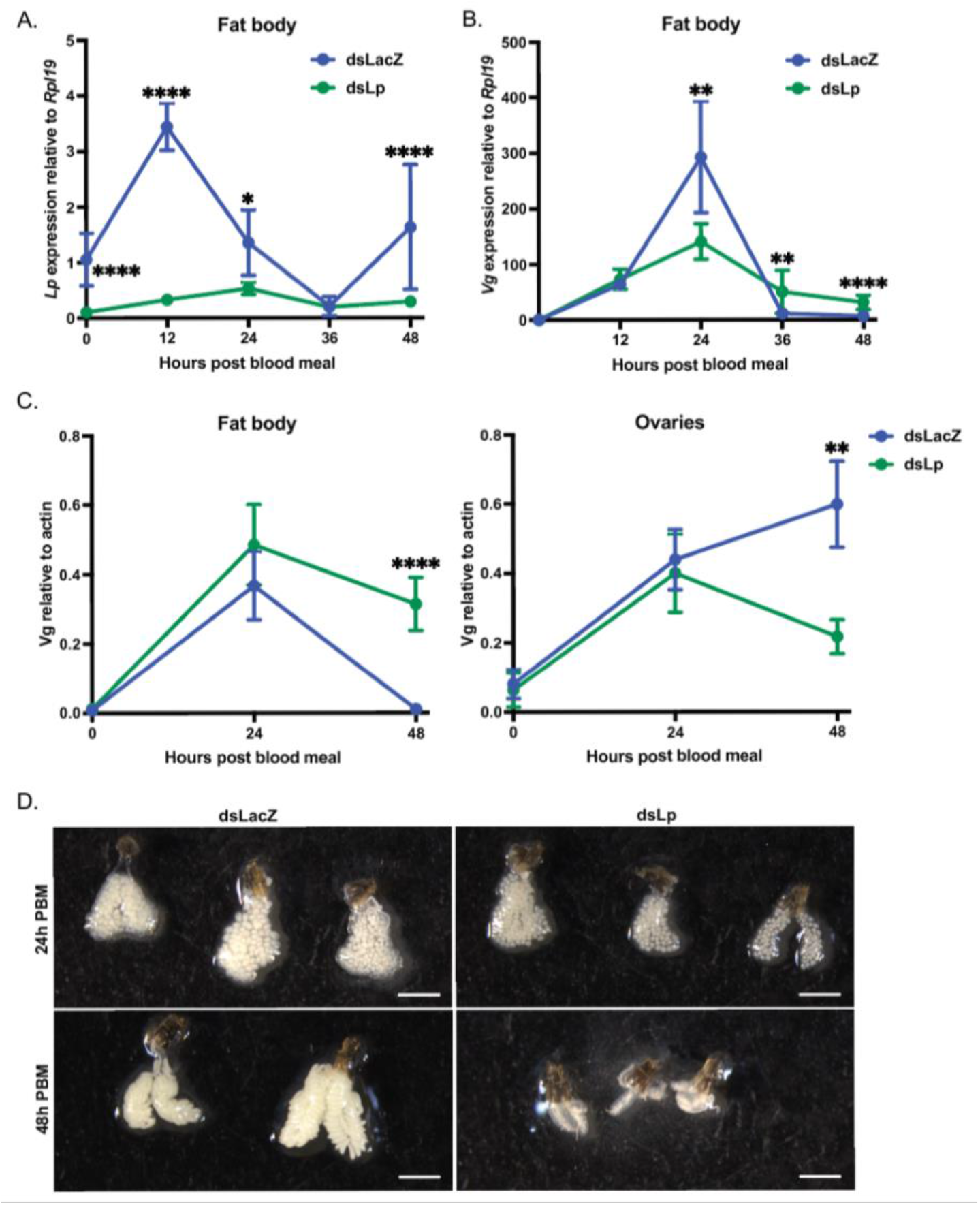
L*p* knockdown significantly impairs oogenesis and affects Vg expression and localization. (A) Successful *Lp* knockdown as determined by RT-qPCR of *Lp* expression levels relative to *Rpl19* in the fat body of dsLacZ and dsLp females (REML variance component analysis: * = p < 0.05; **** = p < 0.0001; three biological replicates). (B) RT-qPCR of *Vg* expression levels relative to *Rpl19* in the fat body of dsLacZ and dsLp females (REML variance component analysis: ** = p < 0.01; **** = p < 0.0001; four biological replicates). (C) Western blot quantification from Figure 1C showing an accumulation of Vg in the fat body and a decrease of Vg in the ovaries upon *Lp* knockdown (REML variance component analysis: ** = p < 0.01; **** = p < 0.0001; three biological replicates). (D) Images of ovaries at 24 and 48h post blood meal showing that Lp depleted ovaries develop normally at first before degenerating by 48h; scale bar = 2mm.

### *Vg* silencing upregulates *Lp* expression and enhances lipid deposition into oocytes

We next assessed the role of *Vg* in egg development and fertility by silencing this gene in females prior to blood feeding (Supplemental Figure 2A). As expected, given its main role during vitellogenesis, Vg depletion caused a significant reduction in number of eggs compared to control females (Figure 2A). More strikingly, however, silencing of this yolk gene induced complete infertility, with no embryos hatching from Vg-depleted mothers (Figure 2B). To confirm this phenotype, we tested a second dsRNA construct targeting *Vg*, and we again observed complete infertility (Figure 2B). No yolk bodies could be detected in dsVg ovaries by electron microscopy (Figure 2C), and we also observed a notable decrease in total protein (Supplemental Figure 2B) and free amino acids (Supplemental Figure 2C) levels, consistent with Vg being a major amino acid source.

**Figure 2.**
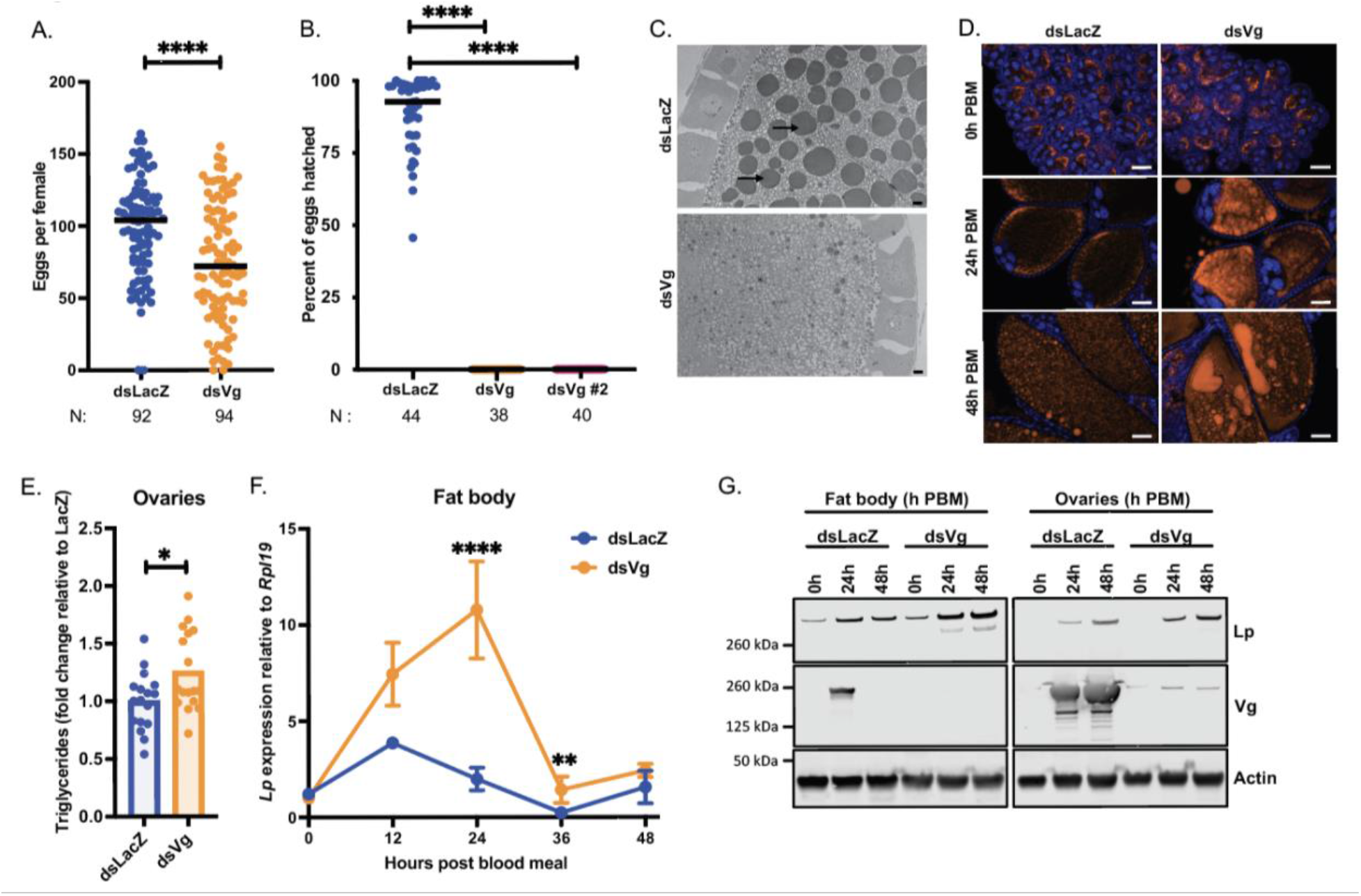
*Vg* silencing upregulates *Lp* expression and enhances lipid deposition into oocytes. (A) Following *Vg* knockdown females develop fewer eggs; each dot represents eggs per female; N = number of females (Mann-Whitney: **** = p < 0.0001; five biological replicates). (B) *Vg* knockdown causes complete infertility; *Vg* was targeted by two different dsRNAs; each dot represents percent hatch rate per female; N = number of females (Kruskal-Wallis with Dunn’s multiple comparisons test: **** = p < 0.0001; three biological replicates). (C) Transmission electron microscopy showing a lack of yolk granules (arrows) upon *Vg* knockdown at 24h post blood meal; scale bar = 2µm (two biological replicates). (D) Fluorescent microscopy showing an accumulation of neutral lipids (LD540, orange) upon *Vg* knockdown; DNA (DAPI) in blue; scale bar = 50µm (three biological replicates). (E) Triglyceride levels measured in dsLacZ and dsVg ovaries at 48h post blood meal and normalized to mean dsLacZ levels in that replicate; each dot is representative of ovaries from three females (Unpaired t-test: * = p < 0.05; three biological replicates). (F-G) *Vg* knockdown results in an increase in Lp levels in the fat body at the mRNA (REML variance component analysis: ** = p < 0.01; **** = p < 0.0001; four biological replicates) (F) and protein (three biological replicates) (G) levels in the fat body and ovaries.

Upon microscopic analysis, we noticed that Vg-depleted ovaries had a remarkable accumulation of neutral lipids at both time points analyzed (24 and 48h PBM) (Figure 2D), a finding which was supported by an assay showing higher triglyceride levels (Figure 2E). Since transport of neutral lipids is mediated by Lp, we assayed *Lp* expression and found that this lipid transporter was upregulated at both transcript and protein levels (Figure 2F, G; Supplemental Figure 2D, E). Specifically, while *Lp* transcripts in the fat body (the tissue where Lp is synthetized) peaked at 12h PBM in controls, after *Vg* silencing they peaked later and at higher levels relative to controls (Figure 2F, Supplemental Figure 2D), paralleled by higher protein levels in the fat body and the ovaries (Figure 2G, Supplemental Figure 2E). Combined, these data suggest that Vg synthesis in the fat body and/or its incorporation into the ovaries is a signal that regulates *Lp* expression and prevents excessive Lp-mediated lipid accumulation in the ovaries, ensuring correct balance between nutrients and safeguarding fertility.

**Supplemental Figure 2.**
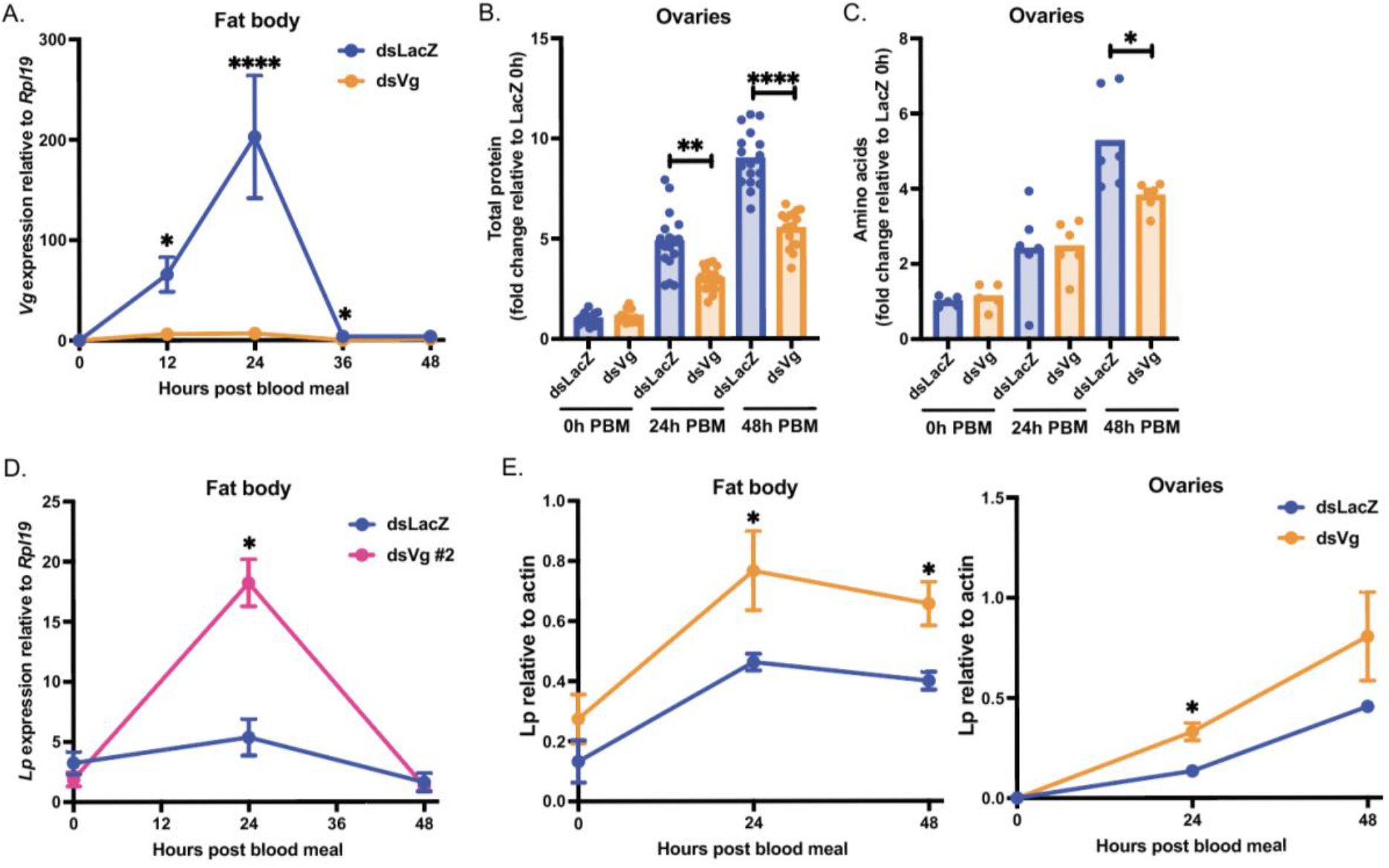
V*g* knockdown effects in fat body and ovaries. (A) Successful *Vg* knockdown as determined by RT-qPCR of *Vg* expression levels relative to *Rpl19* in the fat body (REML variance component analysis: * = p < 0.05; **** = p < 0.0001; three biological replicates). (B) Fold change in protein levels measured by Bradford assay in the ovaries of dsLacZ and dsVg females before blood meal and at 24h and 48h post blood meal (PBM); each dot is representative of three ovaries (REML variance component analysis by timepoint: ** = p < 0.01; **** = p < 0.0001; three biological replicates). (C) Fold change in free amino acid levels in the ovaries of dsLacZ and dsVg females before blood meal and at 24h and 48h PBM; each dot is representative of five ovaries (REML variance component analysis by timepoint: * = p < 0.05; three biological replicates). (D) *Vg* knockdown by second target also results in an increase in *Lp* levels as determined by RT-qPCR (REML variance component analysis: * = p < 0.05; three biological replicates). (E) Western blot quantification from Figure 2G showing an accumulation of Lp in the fat body and ovaries upon *Vg* knockdown (REML variance component analysis: fat body – dsRNA: p < 0.01; * = p < 0.05; ovaries – dsRNA: p < 0.05; * = p < 0.05; three biological replicates).

### *Vg* expression regulates Lp-mediated accumulation of lipids via TOR signaling

In *Ae. aegypti* mosquitoes, *Vg* transcription is partly regulated by TOR signalling (Attardo, Hansen, and Raikhel 2005; Hansen et al. 2014). As mentioned above, upon sensing amino acids derived from the blood meal, TOR phosphorylates S6K, which in turn activates AaGATAa, a transcription factor that binds the *Vg* promoter. We confirmed that *Vg* is regulated by TOR in *An. gambiae* by using the TOR inhibitor rapamycin, which decreased *Vg* expression in the fat body by almost 50% when applied to females at 2h post PBM (Figure 3A). Surprisingly, we also noticed that *Vg* silencing in turn affected TOR signaling in the fat body, mediating its increase relative to controls. Indeed, while in controls S6K phosphorylation had waned by 24h PBM, *Vg*-silenced females showed a strong signal at this time point, which corresponds to the peak of *Vg* expression (Figure 3B, Supplemental Figure 3A). This was paralleled by significantly higher total protein levels after blood feeding (Supplemental Figure 3B), indicating potential translational upregulation by TOR.

**Figure 3.**
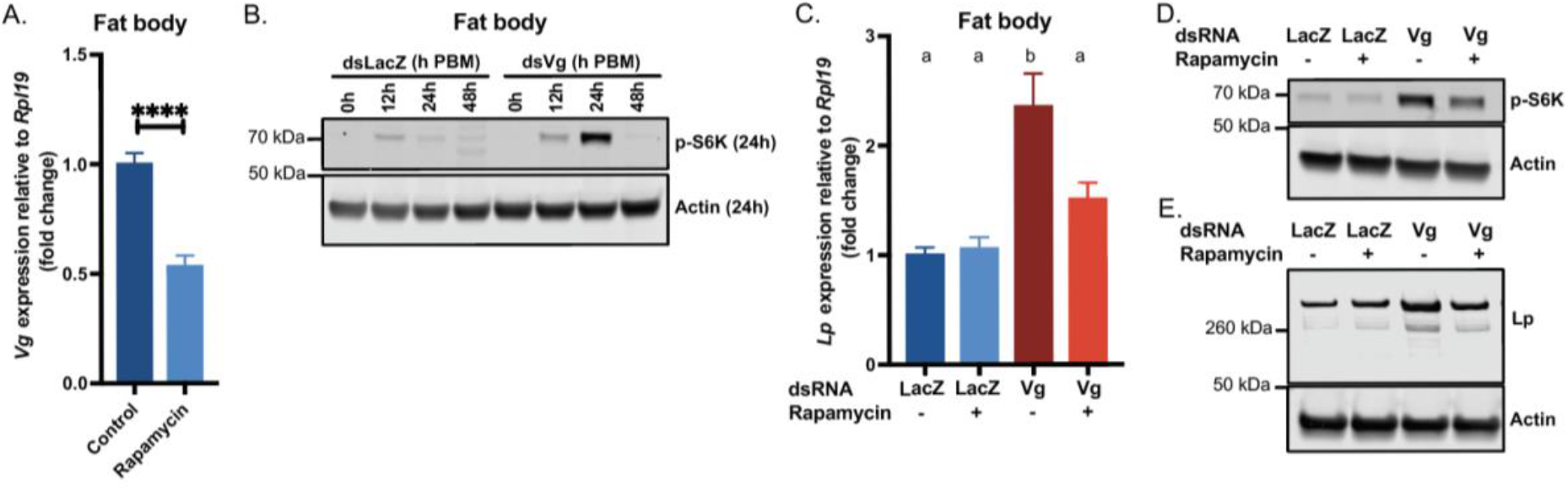
*Vg* regulates *Lp* levels via TOR signaling. (A) *Vg* expression relative to *Rpl19* is reduced in the fat body 24h PBM upon 0.5 µl of 40µM rapamycin treatment 2h PBM; Control is 2.4% DMSO in acetone; all values normalized to control (unpaired t test: **** = p < 0.0001; four biological replicates). (B) TOR signaling is induced in the fat body upon *Vg* knockdown as shown by Western blotting of phospho-S6K levels (three biological replicates). (C) The *Vg*-mediated upregulation of *Lp* at 24h PBM is suppressed in the fat body of ds*Vg* females treated with 0.5 µl of 40 µM rapamycin 2h PBM; all values normalized to ds*LacZ* without rapamycin treatment (ANOVA; four biological replicates). (D) Western blot showing phospho-S6K levels increase in the fat body at 24h PBM in *Vg*-silenced females, but this increase is repressed by rapamycin treatment (four biological replicates). (E) Western blot showing fat body Lp levels at 48h PBM in the same four groups (four biological replicates).

To determine whether TOR activation may be involved in the increased *Lp* expression observed in *Vg*-silenced females, we treated Vg-depleted mosquitoes with rapamycin and assessed *Lp* levels relative to controls. Rapamycin treatment 2h PBM reduced S6K phosphorylation at 24h PBM following Vg depletion, confirming that this phosphorylation is indeed mediated by TOR (Figure 3D, Supplemental Figure 3C). *Lp* upregulation was also significantly reduced (at both protein and transcript levels) by rapamycin treatment, thereby implicating TOR signaling in the Vg-mediated regulation of this lipid transporter (Figure 3C, E, Supplemental Figure 3D). In agreement with this observation, rapamycin treatment also slightly reduced the excess lipid accumulation observed upon *Vg* depletion (Supplemental Figure 3E). Overall, these data suggest that in normal conditions *Vg* expression represses TOR signaling, putting a break on Lp synthesis thereby preventing excessive incorporation of lipids into the ovaries.

We hypothesized that excess amino acids not incorporated into Vg may be the triggers for activating TOR signaling. Indeed, in anopheline mosquitoes, yolk production utilizes up to 30% of the total protein content from the blood meal (Briegel 1990). Consistent with this hypothesis, we detected a modest but significant increase in amino acid levels at 24h PBM in the fat body (but not in the hemolymph) of *Vg*-silenced females, which was amplified at 48h PBM (Supplemental Figure 3F, G). TOR also incorporates inputs from ILPs to regulate metabolism (Arsic and Guerin 2008), however expression levels of 7 *An. gambiae ILPs* were similar between groups at all time points analyzed (Supplemental Figure 3H), likely ruling out insulin signaling as the cause of increased TOR signaling (note, ILP1 and 7, and ILP 3 and 6 share sequence identity, resulting in 5 RT-qPCR plots).

Based on these data, a model emerges whereby in *An. gambiae* TOR signaling both controls *Vg* expression and is in turn regulated by this yolk protein, affecting the expression of *Lp* and the timely accumulation of lipids in the developing ovaries (see graphical abstract). Fine tuning of the function of these nutrient transporters is essential to support egg development and ensure fertility, demonstrating a previously unappreciated interplay that is key to the survival of this species.

**Supplemental Figure 3.**
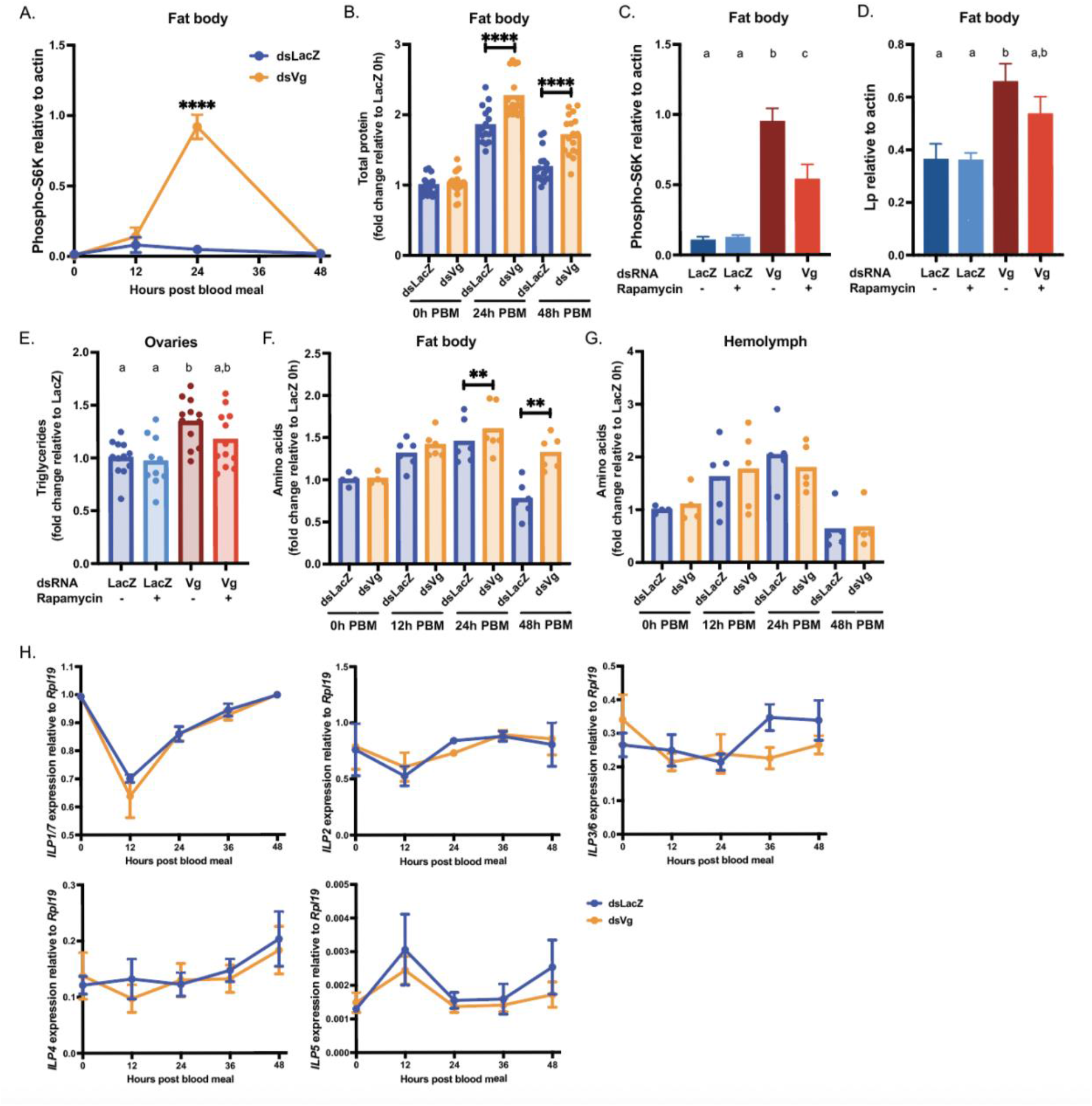
V*g* expression regulates Lp-mediated accumulation of lipids via TOR signaling. (A) Western blot quantification from Figure 3B showing an increase in phospho-S6K levels in the fat body upon *Vg* knockdown (REML variance component analysis: **** = p < 0.0001; three biological replicates). (B) Fold change in protein levels, measured by Bradford assay of the fat body are increased in ds*Vg* females; each dot is representative of three fat bodies (REML variance component analysis by timepoint: **** = p < 0.0001; three biological replicates). (C) Western blot quantification from Figure 3D showing a decrease in phospho-S6K levels in the fat body upon rapamycin treatment (ANOVA; three biological replicates). (D) Western blot quantification from Figure 3E showing Lp protein levels upon *Vg* knockdown and rapamycin treatment (ANOVA; three biological replicates). (E) Triglyceride levels measured in dsLacZ and dsVg ovaries upon 0.5 µl of 40µM rapamycin treatment at 72h post blood meal and normalized to mean dsLacZ levels in that replicate; each dot is representative of ovaries from three females (ANOVA; two biological replicates). (F) Fold change in free amino acid levels in the fat bodies of dsLacZ and dsVg females before blood meal and at 12h, 24h and 48h post blood meal; each dot is representative of five ovaries (REML variance component analysis by timepoint: ** = p < 0.01; three biological replicates). (G) Fold change in free amino acid levels in the hemolymph of dsLacZ and dsVg females before blood meal and at 12h, 24h and 48h post blood meal; each dot is representative of hemolymphs collected from five females (REML variance component analysis by timepoint; three biological replicates). (H) RT-qPCR of *ILP* expression levels relative to *Rpl19* in the heads of dsLacZ and dsVg females (REML variance component analysis; ILP1/7: two biological replicates; other ILPs: three biological replicates).

### *Vg* knockdown causes early embryonic arrest

Our discovery that close cooperation between Lp and Vg is fundamental to fertility prompted us to dig deeper into the mechanisms causing infertility in embryos laid by Vg-depleted females. We found that most of these embryos were fertilized but not melanized (Supplemental Figure 4A) and did not reach the blastocyst stage, which control embryos instead reached at 3-5h post oviposition (Figure 4A). Some embryos were halted very early in development (Figure 4A, top panel), upon the first few mitotic divisions after the fusion of the gamete nuclei, while others reached the energid stage, where nuclei have divided and begin to migrate toward the outer edges (Figure 4A, bottom panel and zoomed-in insert). However, these nuclei displayed abnormal morphology, showing features reminiscent of apoptotic blebbing.

**Figure 4.**
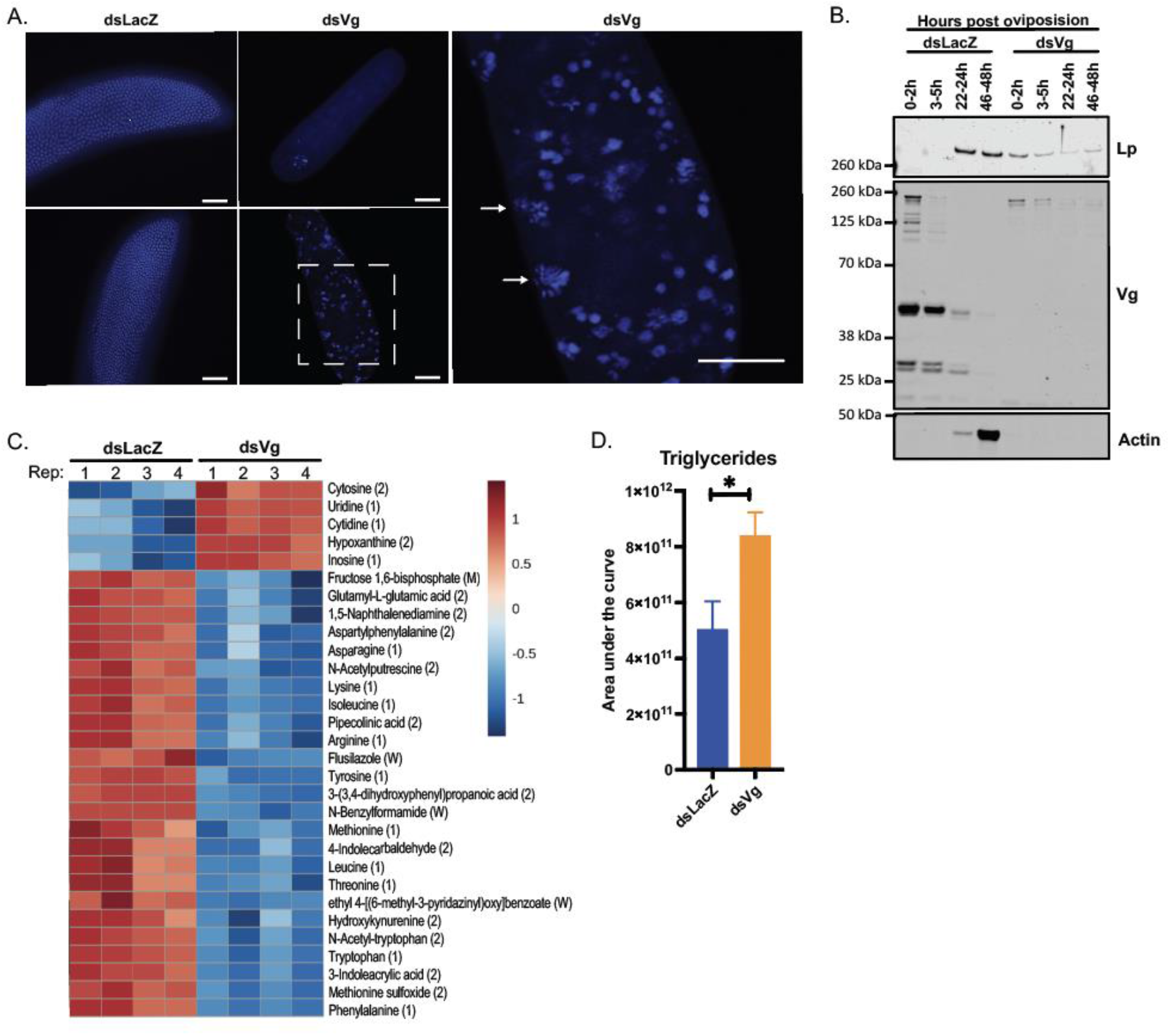
*Vg* knockdown causes early embryonic arrest likely due to limited amino acids. (A) DAPI staining of embryos from ds*LacZ* and ds*Vg* females at 3-5h post oviposition showing arrest of development before the cellular blastocyst stage in the ds*Vg*-derived embryos; scale bar = 50µm (two biological replicates); dotted box represents the field of view in the zoomed-in image to the right, and arrows point to nuclei characteristic of blebbing. (B) Lp is deposited into ds*Vg*-derived embryos and not ds*LacZ*-derived ones, as observed by Western blot at specified time post oviposition (three biological replicates). (C) Top 30 dysregulated metabolites (as determined by Metaboanalyst), of which 10 are amino acids, in embryos derived from ds*Vg* mothers compared to those of ds*LacZ* 3-5h post oviposition (each column is representative of a biological replicate; four biological replicates). ID confidence is in brackets; from strongest to weakest: (1) = Level 1 ID, (2) = Level 2 ID, (M) = MasslistRT ID, (W) = Weak/Poor ID. (D) Triglyceride levels are elevated in ds*Vg*-derived embryos 3-5h post oviposition as determined by lipidomics (unpaired t test: * = p < 0.05; four biological replicates).

We did not detect any Vg in embryos deposited by Vg-depleted mothers, contrary to controls and demonstrating that in early stages of embryogenesis Vg is entirely maternally derived (Figure 4B). Interestingly, we detected Lp in those embryos at very early stages (0-2h) post oviposition, suggestive of maternal deposition which however did not occur in controls (Figure 4B). Moreover, dsVg-derived embryos showed significantly reduced protein synthesis, as highlighted by undetectable actin expression over time (Figure 4B).

In agreement with these findings, metabolomic analysis of dsVg-derived embryos (3-5 hours post oviposition) showed a depletion of amino acids, especially severe for those amino acids that are most prevalent in Vg (Figure 4C; Supplemental Tables 1, 2). Intriguingly, the two amino acids more strikingly depleted, phenylalanine and tyrosine, are the starting compounds in the melanization pathway (Kim et al. 2005; González-Santoyo and Córdoba-Aguilar 2012), possibly explaining the lack of chorion darkening observed in dsVg embryos (Supplemental Figure 4A). Also notable was the upregulation of free nucleotides as well as products of their catabolism such as hypoxanthine and inosine, likely due to impaired DNA replication leading to cell death (Figure 4C).

Given the observed incorporation of Lp into the embryo, we also performed lipidomic analysis at the same time point. Although there were no differences in total lipids in dsVg-derived embryos relative to controls (Supplemental Figures 4B, Supplemental Table 3), some minor lipid classes (phospholipids and glycerophospholipids) were decreased (Supplemental Figure 4C), while we detected a remarkable increase in triglycerides (Figure 4D), again consistent with the known role of Lp in transport of these lipids (Ford and Van Heusden 1994).

Therefore, when mothers are depleted of Vg, embryos cannot develop further than the first steps after fertilization, likely due to a severe drop in amino acid levels and protein synthesis.

**Supplemental Table 1:** Vg amino acids and their decrease in embryos upon *Vg* depletion.

| **Amino acid** | **Percent content in Vg** | **Decrease upon Vg KD in mothers (fold change)** | **Significance** |  |
| --- | --- | --- | --- | --- |
| Ser (S) | 8.50% | 2.2 | 0.09205 |  |
| Tyr (Y) | 8.00% | 52.6 | 0.00010 | *** |
| Phe (F) | 7.70% | 63.2 | 0.00000 | **** |
| Glu (E) | 6.50% | 1.2 | 0.33025 |  |
| Ala (A) | 6.30% | 2.0 | 0.02859 | * |
| Asp (D) | 6.30% | 2.1 | 0.15012 |  |
| Lys (K) | 6.20% | 10.0 | 0.00000 | **** |
| Gln (Q) | 6.10% | 1.1 | 0.88614 |  |
| Val (V) | 5.80% | 3.1 | 0.00005 | **** |
| Asn (N) | 5.60% | 5.5 | 0.00006 | **** |
| Leu (L) | 5.60% | 3.7 | 0.00024 | *** |
| Gly (G) | 4.40% | 2.2 | 0.02060 | * |
| Pro (P) | 4.40% | 1.4 | 0.01066 | * |
| Thr (T) | 4.40% | 3.9 | 0.00015 | *** |
| Arg (R) | 3.90% | 2.9 | 0.00012 | *** |
| Ile (I) | 3.20% | 8.7 | 0.00001 | **** |
| His (H) | 2.90% | 1.0 | 0.69815 |  |
| Met (M) | 2.20% | 4.9 | 0.00010 | *** |
| Cys (C) | 1.10% | Not detected |  |  |
| Trp (W) | 0.80% | 13.2 | 0.00002 | **** |

**Supplemental Figure 4.**
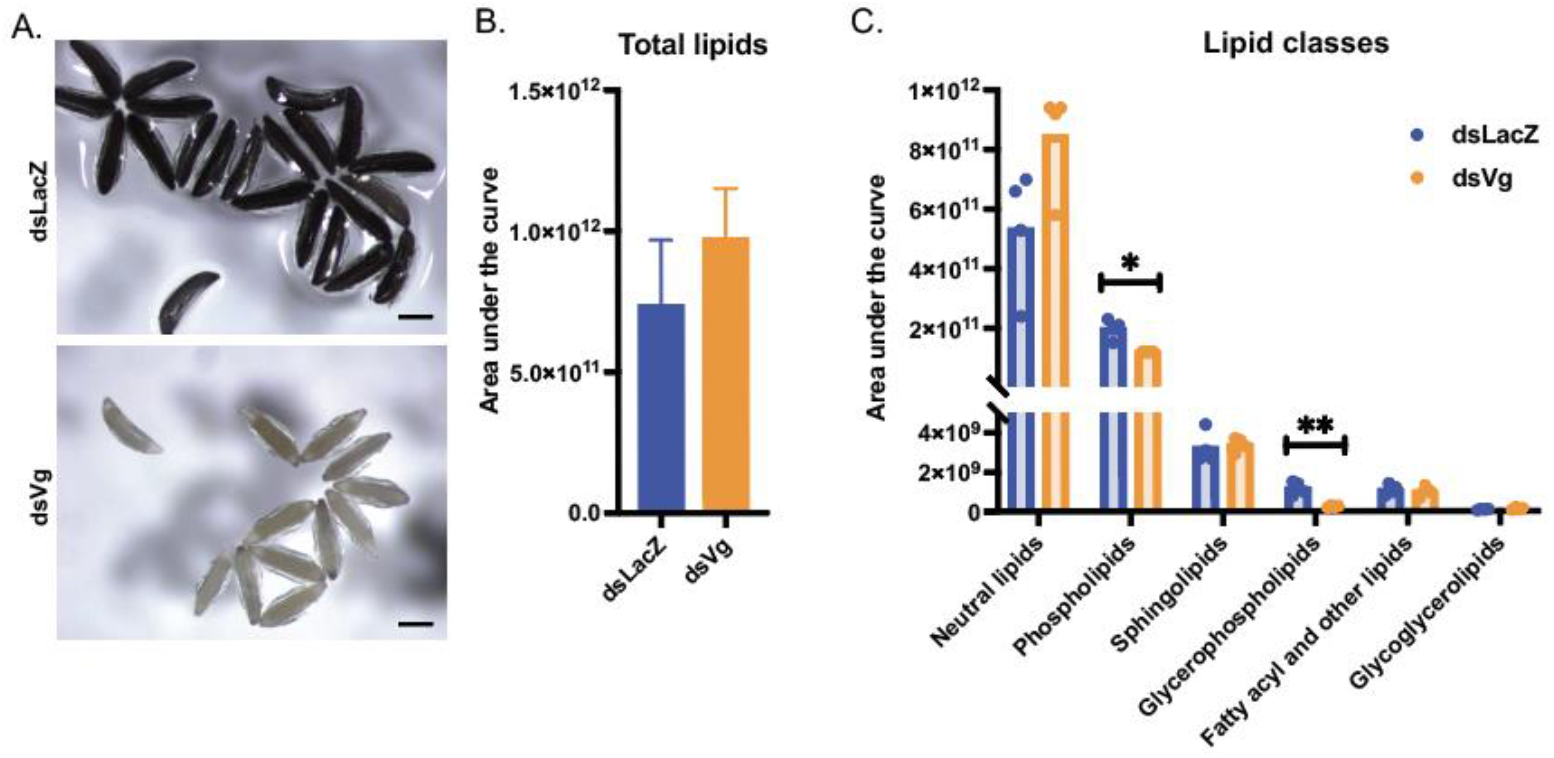
V*g* knockdown in females prevents embryo melanization and causes early embryonic arrest. (A) Light microscopy of embryos from dsLacZ- and dsVg-derived females at 3-5h post oviposition; scale bar = 200 µm. (B-C) Total lipids (B) and lipid classes (C) in dsLacZ- and dsVg-derived embryos 3-5h post oviposition as determined by mass spectrometry (unpaired t tests, followed by FDR correction: * = p < 0.05; ** = p < 0.01; four biological replicates).

## Discussion

Blood feeding is an essential process for the survival of mosquito species that, like *An. gambiae*, rely on blood nutrients for oogenesis. After a blood meal, different nutrient transporters start shuffling cholesterol, lipids and proteins from the midgut and fat body to the ovaries, and the individual roles of two of these factors, Lp and Vg, had been previously determined. There remained, however, important outstanding questions concerning whether and how these two essential transporters are co-regulated in order to ensure reproductive success. How do mosquitoes balance acquisition of different nutrients by the ovaries? What are the signals that limit lipid accumulation and that trigger vitellogenesis? Moreover, knowledge concerning Lp and Vg functions and their regulation in *Anopheles* as opposed to *Aedes* was only marginal. Here, we show that in these mosquitoes, oogenesis is the product of a precise and intricate interplay between these two factors. The initial phase of egg development is dominated by Lp, which incorporates lipids (mostly triglycerides) and cholesterol into ovaries triggering their growth. During this early phase, as 20E is synthesized in the fat body, this steroid hormone triggers the expression of *Vg*, which peaks at 24h PBM and provides the oocytes with considerable stores of amino acids. When *Lp* expression is impaired, egg development is strikingly reduced (Figure 1A), Vg localization is affected (Figure 1C), and yolk bodies are aberrant (Figure 1D). These findings suggest that correct lipid accumulation is an early check point that mosquitoes use to decide whether to proceed with vitellogenesis. Reducing *Vg* expression levels, on the other hand, leads to increased *Lp* expression (Figure 2F and G) and an overload of glycerides in the ovaries (Figure 2E), which suggests that Vg synthesis is the signal that prevents excessive lipid trafficking by Lp into the oocytes. Interestingly, in another study *Vg* silencing did not appear to affect *Lp* expression in *An. gambiae* females fed on mice infected with *P. berghei* parasites (Rono et al. 2010). This discrepancy with our results may indicate that rodent malaria parasites, which are known to inflict severe reproductive costs in infected mosquito females (Ahmed and Hurd 2006), affect the normal accumulation of lipids during the oogenetic process, although other factors such as differences in temperature (infections with *P. berghei* are done at permissive temperatures around 20°C compared to standard mosquito rearing conditions of 28°C) cannot be excluded.

Strikingly, silencing of *Vg* led to complete infertility. This phenotype was so penetrant that we confirmed it with a dsRNA second construct targeting a different region of the gene to rule out a possible unspecific effect of the first construct. This effect is reminiscent of observations in other insects, where altering Vg gene copy number, expression or internalization leads to complete sterility or decreased hatch rate by as yet unknown mechanisms (Bownes, Lineruth, and Mauchline 1991; DiMario and Mahowald 1987; Gutzeit and Arendt 1994; Lin et al. 2013; Ciudad, Piulachs, and Bellés 2006; Peng et al. 2020; Shang et al. 2018; Mitchell III et al. 2007; Coelho et al. 2016). Understanding how embryonic lethality is induced may lead to novel ideas for the design of mosquito-targeted interventions, so we set out to determine the mechanisms behind death. As *Vg* is also expressed in the female spermatheca after mating (Rogers et al. 2008; Shaw et al. 2014) we initially thought that its depletion may have caused irreversible damage to sperm. The observation that embryos are fertilized and start undergoing nuclear division (Figure 4A), however, appears to discount this possibility. Based on our metabolomics analysis, the most plausible hypothesis is that infertility is a result of amino acid starvation. We show that embryos from Vg-depleted females are significantly deficient in 14 of the 19 identified amino acids, which results in a lack of building blocks for translation and thus development (Supplemental Table 1). It is plausible to speculate that depletion in these essential nutrients may activate the amino acid response pathway, triggering a global shutdown of translation that may lead to apoptosis, compatible with our observation of nuclei blebbing in those embryos (Sikalidis 2013; Dong et al. 2000; Qin et al. 2017).

Our data using rapamycin suggest that the *An. gambiae* Vg is regulated by TOR (Figure 3A), as was previously shown in *Ae. aegypti* (Hansen et al. 2004). Surprisingly, however, our findings also suggest that Vg in turn suppresses TOR signaling, as upon *Vg* knockdown S6K phosphorylation was strongly upregulated possibly due to an increase in free amino acids (Figure 3B and Supplemental Figure 3F). This upregulation in TOR signaling also resulted in an increase in Lp transcription and translation, further shedding light on regulation of *Lp* expression. Previous studies had shown that *Lp* levels in *An. gambiae* are under steroid hormone control, as impairing 20E signaling caused an increase in *Lp* transcription after blood feeding (Werling et al. 2019). With the caveat that our current results were obtained only by using the inhibitor rapamycin rather than by also silencing key components of the TOR pathway, these combined observations may suggest that TOR and 20E signaling exert opposite effects on *Lp* expression — with 20E repressing its levels and TOR enhancing them —, an intriguing finding that deserves more thorough investigation in future studies. Compatible with our data, the Lp promoter has putative GATA transcription factor binding motifs, some of which are known to be regulated by TOR signaling (Marinotti et al. 2006; Park et al. 2006).

Does the interplay between Lp and Vg also affect the development of *P. falciparum* parasites? An earlier study showed that when *Lp* is silenced in *An. gambiae*, *P. falciparum* oocyst numbers are decreased (Werling et al. 2019). No other effects were detected on parasite development, unlike in the mouse malaria parasite *P. berghei* where Lp depletion, besides a decrease in oocyst numbers, also led to reduced oocyst growth (Rono et al. 2010). While the role of Vg in *P. falciparum* has not been directly determined, it is known that impairing 20E signaling (which in turn negatively affects *Vg* levels) has profound and opposite effects on parasites, as it reduces parasite numbers but accelerates their growth. Regardless of its impact on parasite development, our data reveal the interplay between Lp and Vg as essential for mosquito fertility, opening the possibility of targeting it to reduce the reproductive success of mosquito populations.

## Materials and Methods

### Mosquito lines and rearing

G3 *Anopheles gambiae* mosquitoes were reared at 27°C, 70-80% humidity. Adults were fed 10% glucose solution and purchased human blood (Research Blood Components, Boston, MA). Females and males were separated by pupae sexing, and females were kept separate to ensure virgin status or mixed with males at a 1:2 ratio for fertility experiments and egg collections.

### dsRNA generation

A 816bp LacZ fragment and 600bp Lp (AGAP001826) fragment were generated from plasmids pLL100-LacZ and pLL10-Lp as described previously (Blandin et al. 2004; Marcenac et al. 2020; Werling et al. 2019) using T7 primer (5’–TAATACGACTCACTATAGGG–3’). A 552 bp fragment of Vg (AGAP004203) corresponding to bases 3374–3925 of the Vg cDNA was amplified from plasmid pLL10-Vg, a gift from Miranda Whitten and Elena A. Levashina (Max Planck Institute for Infection Biology, Berlin), using a primer matching the inverted T7 promoters (same as above). To generate the dsVg #2 construct, a 284bp PCR product was generated from *An. gambiae* Vg cDNA (AGAP004203) corresponding to bases 4530–4813 using forward primer ATTGGGTACCGGGCCCCCCCGCACGTCTCGATGAAGGGTA and reverse primer GGGCCGCGGTGGCGGCCGCTCTAGACCTGCCCTGGAAGAAGTAGTCC. The pLL10-Vg backbone and the PCR fragment were restriction digested with XbaI and XhoI, separated on an agarose gel and gel purified. Then, fragments were assembled using NEBuilder HiFi DNA Assembly Kit. PCR product was amplified using T7 primer. PCR for dsRNA generation was separated by gel electrophoresis for size confirmation, and transcribed into dsRNA by *in vitro* transcription Megascript T7 kit (ThermoFisher Scientific) (Werling et al. 2019). dsRNA was purified by phenol-chloroform extraction, and diluted to 10µg/µl.

### dsRNA injections

Females on day 1 post eclosion were injected with 69nl of dsRNA (dsLacZ, dsVg, dsVg #2, dsLp) using Nanoject III (Drummond), and allowed to recover. Surviving females were fed with blood 3 days post injection. Unfed females were removed from experimental cages.

### Egg counts

Virgin females were dissected 3-7 d after bloodfeeding, and the egg clutches were counted.

### Fertility assay

Injected females were mixed with males at a 1:2 ratio immediately after injection, and bloodfed three days later. One day after bloodfeeding, fed females were moved to individual cups with around 2cm of water at the bottom. Hatched and unhatched eggs from every cup were counted within a week.

### RNA extraction, cDNA synthesis and RT-qPCR

Fat bodies or heads (10 tissues per tube) from female mosquitoes were dissected in PBS and stored at -80°C in 300µl TRI Reagent (ThermoFisher Scientific). Samples were thawed and bead beaten using 2mm beads. Then RNA was extracted using manufacturer’s instructions with a modification to wash the RNA pellet using 70% ethanol. 2.5µg of RNA was aliquoted and DNase treated with Turbo DNase from the TURBO DNA-free Kit (ThermoFisher Scientific), followed by DNase inactivation from the same kit. cDNA synthesis was carried out in 100µl reactions using random primers (ThermoFisher Scientific), dNTPs (ThermoFisher Scientific), first strand buffer (VWR), RNAseOUT (ThermoFisher Scientific) and MMLV (ThermoFisher Scientific). Relative quantification RT-qPCR was carried out using SYBR-Green mix and primers from Supplemental Table 4. Primers were designed on exon-exon junctions where possible. Quantification was performed in triplicate using the QuantStudio 6 Pro qPCR machine (ThermoFisher Scientific). Rpl19 was used as the endogenous control for relative quantification.

**Supplemental Table 4.**
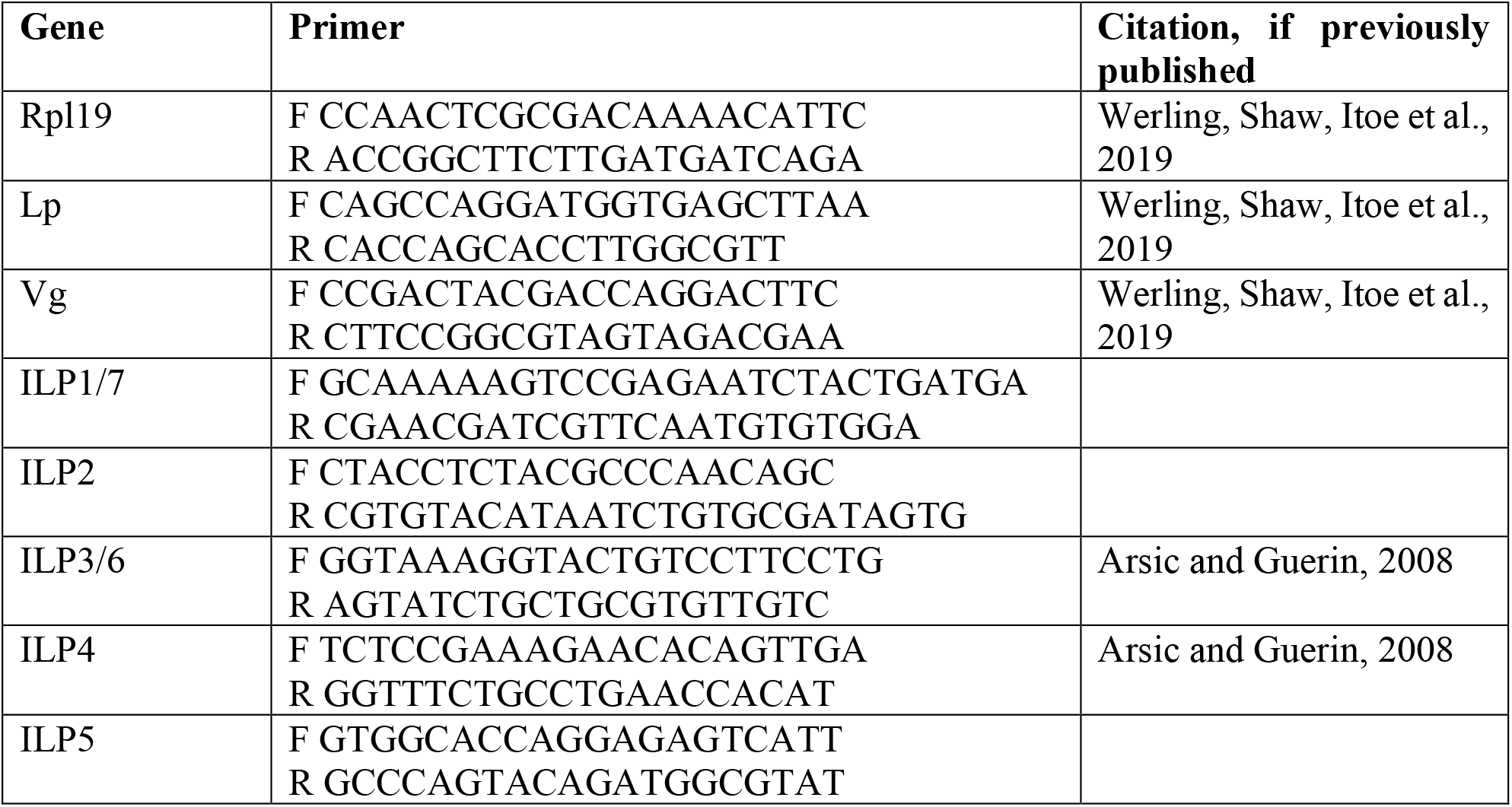

### Immunofluorescent microscopy and tissue staining

Ovaries were dissected from females at specified timepoints and incubated at room temperature in 4% paraformaldehyde (PFA) for 30 minutes, followed by 3 washes in PBS for 15 minutes. Ovaries were permeabilized and blocked with 0.1% Triton X-100 and 1% bovine serum albumin (BSA) in PBS for 1 hour followed by 3 washes in PBS for 15 minutes. Ovaries were then incubated with DAPI and LD540 (Spandl et al. 2009), both at a concentration of 1µg/ml, at room temperature for 15 minutes. After staining, ovaries were washed in PBS 3 times for 15 minutes and mounted using Vectashield mounting medium (Vector Laboratories). Images were captured on a Zeiss Inverted Observer Z1.

### Embryo collections for microscopy

An egg bowl was inserted into cages of mated females bloodfed 96h before. The egg bowl was removed 2 hours later, and embryos were collected 3 hours later, resulting in a timepoint of 3-5 hours. Embryos were dechorionated and cracked as described previously (Goltsev et al. 2004). Briefly, embryos were washed with 25% bleach, collected into glass vials with 9% PFA and heptane, and rotated for 25 minutes. PFA was removed and replaced with deionized water twice. Vials were shaken for another 30 minutes. Then water was replaced with boiling water and incubated in a hot water bath for 30 seconds, and immediately replaced with ice cold water. Both water and heptane were removed and replaced with heptane and methanol. Embryos were swirled vigorously to crack the shell, washed 3 times with methanol and collected into methanol. Embryos were then coaxed out of eggshells (Juhn and James 2012), and stained with DAPI as described above.

### Transmission electron microscopy

Dissected mosquito ovaries were collected in 200 uL of fixative (2.5% paraformaldehyde, 5% glutaraldehyde, 0.06% picric acid in 0.2M cacodylate buffer) and spun down briefly to fully submerge the tissues in fixative. Fixed samples were submitted to the Harvard Medical School Electron Microscopy Core. Samples were washed once in 0.1M cacodylate buffer, twice in water, and then postfixed with 1% osmium tetroxide/1.5% potassium ferrocyanide in water for 1 hour. Samples were then washed twice in water followed by once in 50mM maleate buffer pH 5.15 (MB). Next, the samples were incubated for one hour in 1% uranyl acetate in MB, followed by one wash in MB, and two washes in water. Samples were then subjected to dehydration via an increasing ethanol gradient (50%, 70%, 90%, 100%, 100% ethanol) for 10 minutes each. After dehydration, samples were placed in propylene oxide for one hour and then infiltrated overnight in a 1:1 mixture of propylene oxide and TAAB 812 Resin (https://taab.co.uk/, #T022). The following day, samples were embedded in TAAB 812 Resin and polymerized at 60C for 48 hours. Ultrathin sections (roughly 80nm) were cut on a Reichert Ultracut-S microtome, sections were picked up onto copper grids, stained with lead citrate and imaged in a JEOL 1200EX transmission electron microscope equipped with an AMT 2K CCD camera.

### Rapamycin treatment

0.5µl of 40µM rapamycin was applied on ice to the posterior thoraces of females 2h post bloodfeeding. Rapamycin was in 2.4% DMSO in acetone. Control mosquitoes were treated with 0.5µl of 2.4% DMSO in acetone. Mosquitoes were placed into cages with 10% glucose to recover.

### Hemolymph collections

For amino acid assay, hemolymph was collected by making a tear between the last and second-to-last segments on the abdomen, and injecting 2µl of purified deionized water into each mosquito. A drop of liquid was collected from the abdomen with a pipette. Hemolymphs from five females were pooled for amino acid assay.

### Primary antibodies

Antibody against Vg was generated with Genscript by injecting a rabbit with a Vg peptide (QADYDQDFQTADVKC). Rabbit serum was affinity purified to produce a polyclonal antibody used at 1:1000 for Western blotting.

Anti-Lp antibody was also generated with Genscript using the following peptide: FQRDASTKDEKRSGC (Werling et al. 2019). This antibody was used at 1:4000 for Western blotting.

Anti-actin antibody was acquired from Abcam (MAC237) and used at a dilution of 1:4000.

Phospho-S6K antibody was acquired from MilliporeSigma (07-018-I) and used at a dilution of 1:1000.

### Western blotting

5 tissues per sample, or 40 embryos were collected into 55 µl of PBS with protease and phosphatase inhibitors (cOmplete Mini EDTA free protease inhibitor cocktail, Halt phosphatase inhibitor). DTT (200mM) and NuPAGE LDS Sample Buffer were added. Tissues were bead beaten and boiled for 10 minutes. Then, 1/10^th^ of the sample was loaded onto either a NuPAGE 4-12% Bis-Tris gel or NuPAGE 3-8% Tris Acetate gel (when blotting for Lp). Gels were transferred for 10 minutes at 22V using an iBlot2 machine and iBlot2 PVDF Stacks, blocked in Intercept Blocking Buffer (LI-COR) for 1 hour, and then incubated with antibody overnight at 4°C. Membranes were washed with PBS-T 4 times for 5 minutes, and incubated with LI-COR secondary antibodies (Goat anti-Rat 680LT; Donkey anti-Rabbit 800CW) for 1-2 hours. Membranes were washed again with PBS-T 4 times and once in PBS. Membranes were imaged using a LI-COR developer.

### Triglyceride assay and Bradford assay

Three tissues per sample were collected into NP40 Assay Reagent, and Triglyceride Colorimetric Assay was performed according to manufacturer’s instructions (Cayman). Briefly, tissues were homogenized by bead beating in 32µl of the NP40 Assay Reagent, centrifuged at 10,000g for 10 minutes. 10µl of supernatant was added to 150µl of Enzyme Mixture in duplicate and incubated for 30 minutes at 37°C. Absorbance was measured at 530nm.

Of note, the kit releases glycerol from triglycerides and measures glycerol levels, and does not measure triglyceride levels directly.

The same supernatant was also used for Bradford assay. Supernatant was diluted 1 in 10 and 4µl of diluted supernatant was added to 200µl of Bradford reagent (Bio-Rad) at room temperature, and absorbance was recorded at 595nm.

### Amino acid assay

Five tissues per sample were collected into 50µl of Ultrapure Distilled Water (Invitrogen), and amino acid assay was performed according to manufacturer’s instructions (EnzyChrom L-Amino Acid Assay Kit ELAA-100). Briefly, samples were homogenized by bead beating and centrifuged at 10,000g for 15 minutes. 20µl of supernatant was mixed with Working Reagent in duplicate and incubated at room temperature for 60 minutes. Absorbance was recorded at 570nm.

### Sample collection for metabolomics and lipidomics

200 eggs were collected into 1ml of methanol and homogenized by bead beating with 5 2mm glass beads, then transferred to 8ml glass vials. Tubes were then rinsed with 1ml of methanol that was pooled with the homogenized sample, and 4ml of cold chloroform was added to the glass vials, which were then vortexed for 1 minute. 2ml of water was added and glass vials were vortexed for another minute. Vials were then centrifuged for 10 minutes at 3000*g*. The upper aqueous phase was submitted for metabolomics, and lower chloroform phase was submitted for lipidomics to the Harvard Center for Mass Spectrometry.

### Metabolomics mass spectrometry

Samples were dried down under Nitrogen flow and resuspended in 25 µL of acetonitrile 30% in water. Ten microliter of each sample was used to create a pool sample for MS2/MS3 data acquisition. Samples were analyzed by LC-MS on a Vanquish LC coupled to an ID-X MS (Thermofisher Scientific). Five µL of sample was injected on a ZIC-pHILIC peek-coated column (150 mm x 2.1 mm, 5 micron particles, maintained at 40 °C, SigmaAldrich). Buffer A was 20 mM Ammonium Carbonate, 0.1% Ammonium hydroxide in water and Buffer B was Acetonitrile 97% in water. The LC program was as follow: starting at 93% B, to 40% B in 19 min, then to 0% B in 9 min, maintained at 0% B for 5 min, then back to 93% B in 3 min and re-equilibrated at 93% B for 9 min. The flow rate was maintained at 0.15 mL min^-1^, except for the first 30 seconds where the flow rate was uniformly ramped from 0.05 to 0.15 mL min^-1^. Data was acquired on the ID-X in switching polarities at 120000 resolution, with an AGC target of 1e5, and a m/z range of 65 to 1000. MS1 data was acquired in switching polarities for all samples. MS2 and MS3 data was acquired on the pool sample using the AquirX DeepScan function, with 5 reinjections, separately in positive and negative ion mode. Data was analyzed in Compound Discoverer © software (CD,version 3.3 Thermofisher Scientific). Identification was based on MS2/MS3 matching with a local MS2/MS3 mzvault library and corresponding retention time built with pure standards (Level 1 identification), or on mzcloud match (level 2 identification). Compounds where the retention time and the accurate mass matched an available standard, but for which MS2 data was not acquired are labelled MasslistRT matches. Each match was manually inspected.

Metabolomics heatmap was generated using Metaboanalyst 5.0 by log 10 transforming the area under the curve values for metabolites identified as described above.

### Lipidomics mass spectrometry

Samples were dried down under Nitrogen flow and resuspended in 60 µL of chloroform. LC–MS analyses were modified from (Miraldi et al. 2013) and were performed on an Orbitrap QExactive plus (Thermo Scientific) in line with an Ultimate 3000 LC (Thermo Scientific). Each sample was analyzed separately in positive and negative modes, in top 5 automatic data dependent MSMS mode. Twenty µL of sample was injected on a Biobond C4 column (4.6 × 50 mm, 5 μm, Dikma Technologies, coupled with a C4 guard column). Flow rate was set to 100 μl min^−1^ for 5 min with 0% mobile phase B (MB), then switched to 400 μl min^−1^ for 50 min, with a linear gradient of MB from 20% to 100%. The column was then washed at 500 μl min^−1^ for 8 min at 100% MB before being re-equilibrated for 7min at 0% MB and 500 μl min^−1^. For positive mode runs, buffers consisted for mobile phase A (MA) of 5mM ammonium formate, 0.1 % formic acid and 5% methanol in water, and for MB of 5 mM ammonium formate, 0.1% formic acid, 5% water, 35% methanol in Isopropanol. For negative runs, buffers consisted for MA of 0.03% ammonium hydroxide, 5% methanol in water, and for MB of 0.03% ammonium hydroxide, 5% water, 35% methanol in isopropanol. Lipids were identified and integrated using the Lipidsearch © software (version 4.2.27, Mitsui Knowledge Industry, University of Tokyo). Integrations and peak quality were curated manually before exporting and analyzing the data in Microsoft excel.

### Quantification and statistical analyses

All statistical tests were performed in GraphPad Prism 9.0 and JMP 17 Pro statistical software. The number of replicates and statistical tests performed are mentioned in the figure legend. Detailed outputs of statistical models are provided in the supplementary information. Residual Maximal Likelihood (REML) variance components analysis was used by fitting linear mixed models after data transformation to resemble normality. dsRNA treatment and timepoint were included as fixed effects and replicate as a random effect. If transformation was not possible, a generalized linear model was used instead. Interaction terms were removed when not significant and models with lower AICc scores were kept. Multiple comparisons were calculated using pairwise Student’s t tests at each timepoint followed by FDR correction at a 0.05 significance level.

**Supplemental Table 5.**
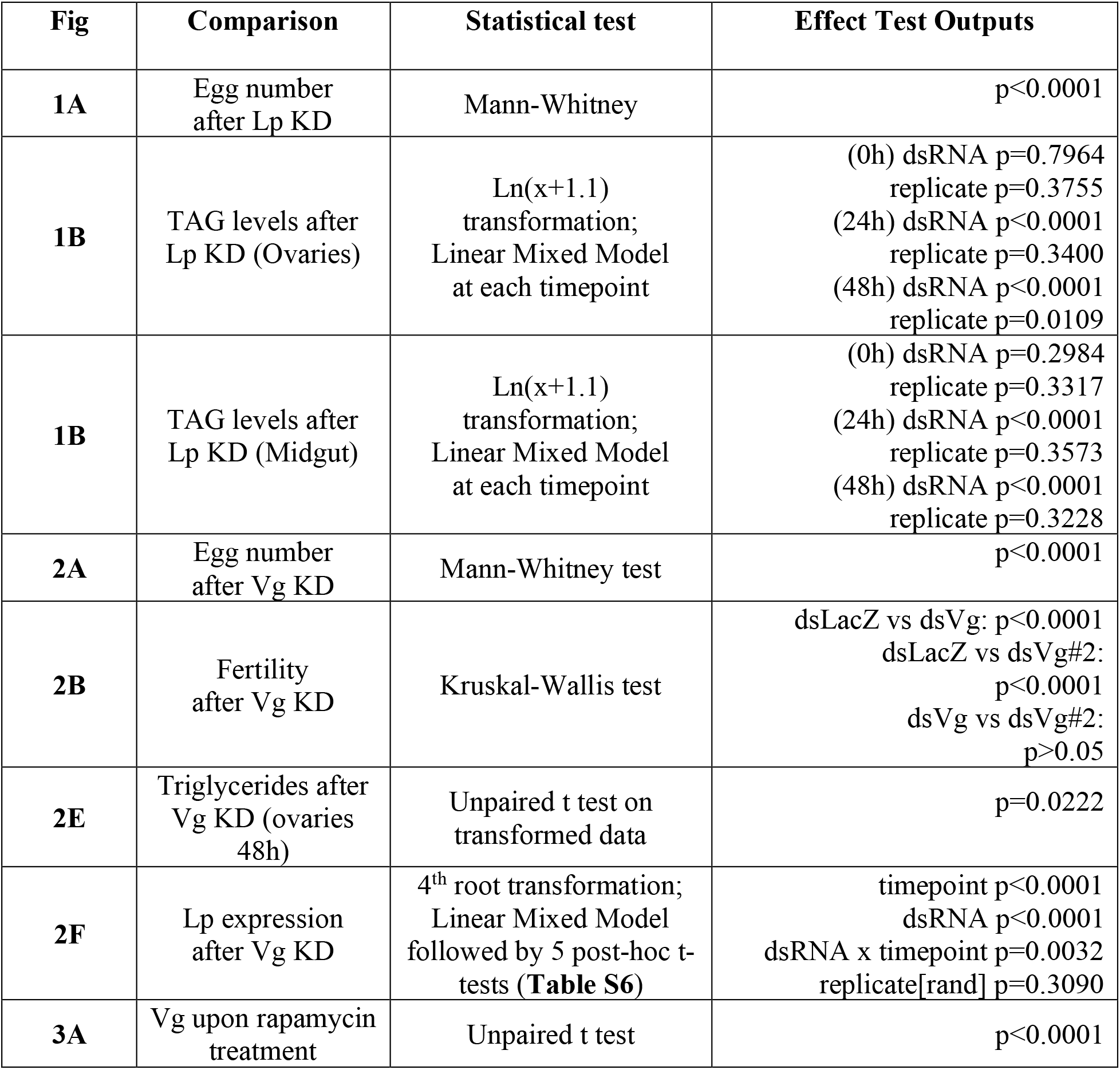
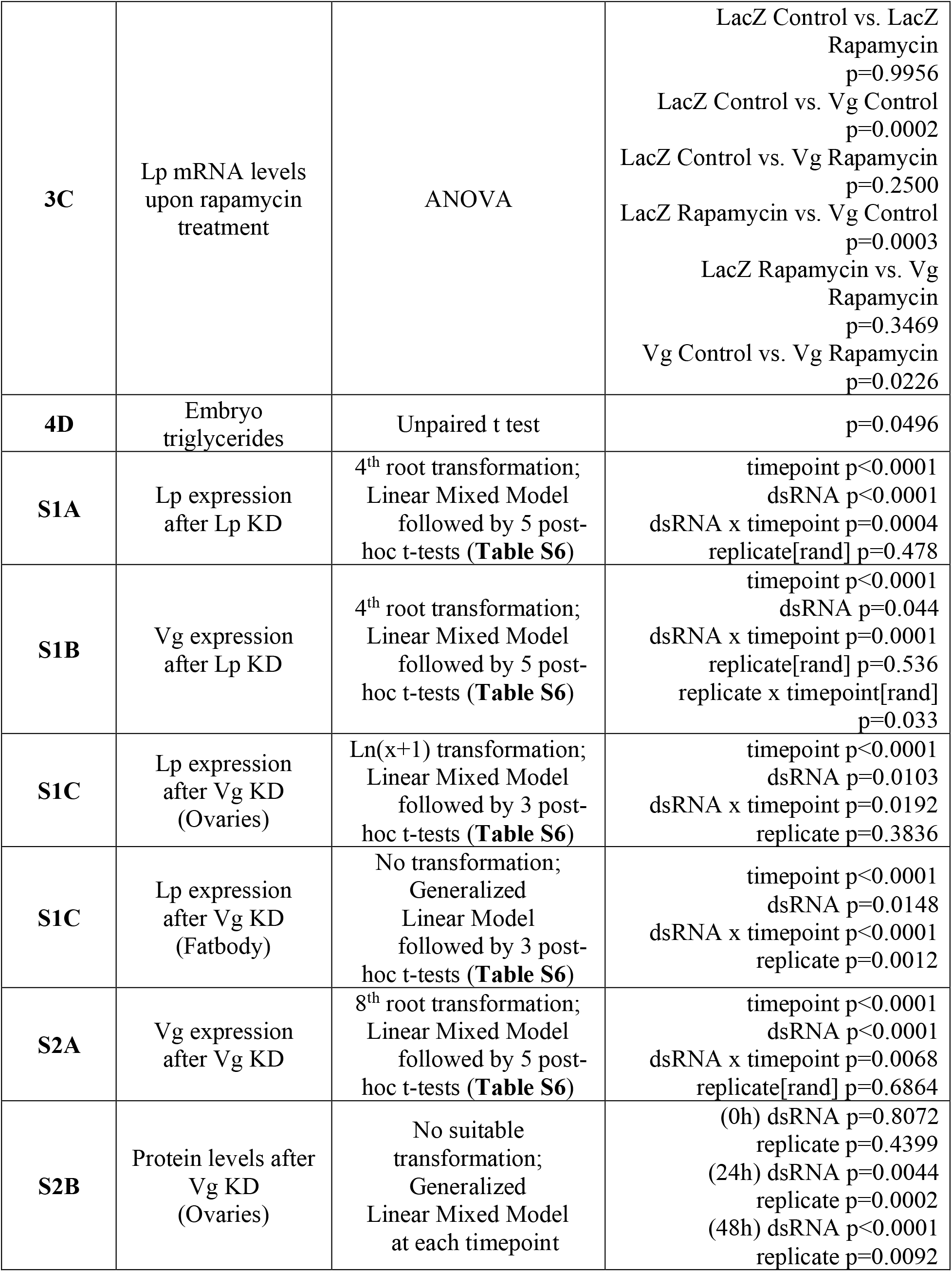
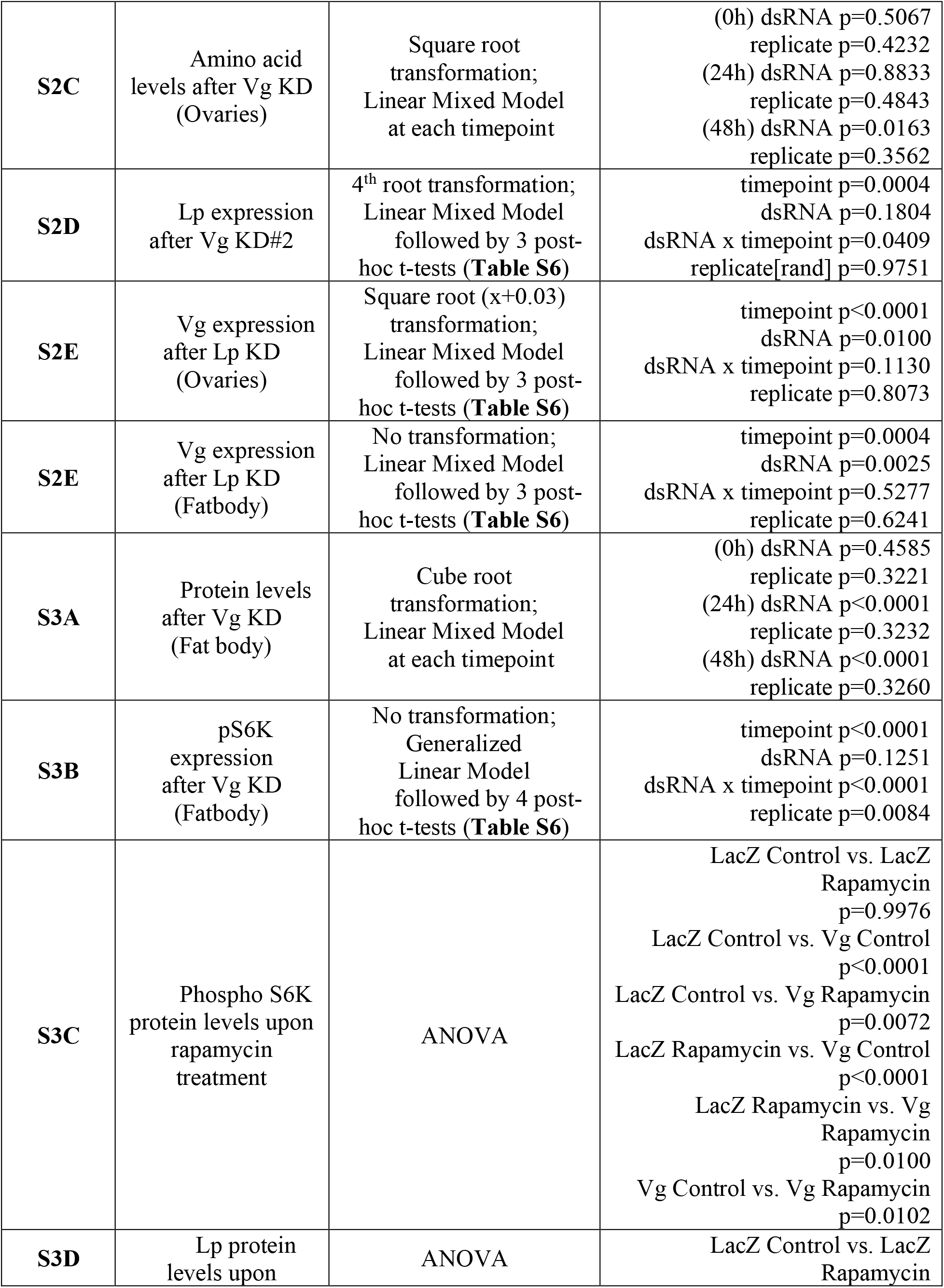
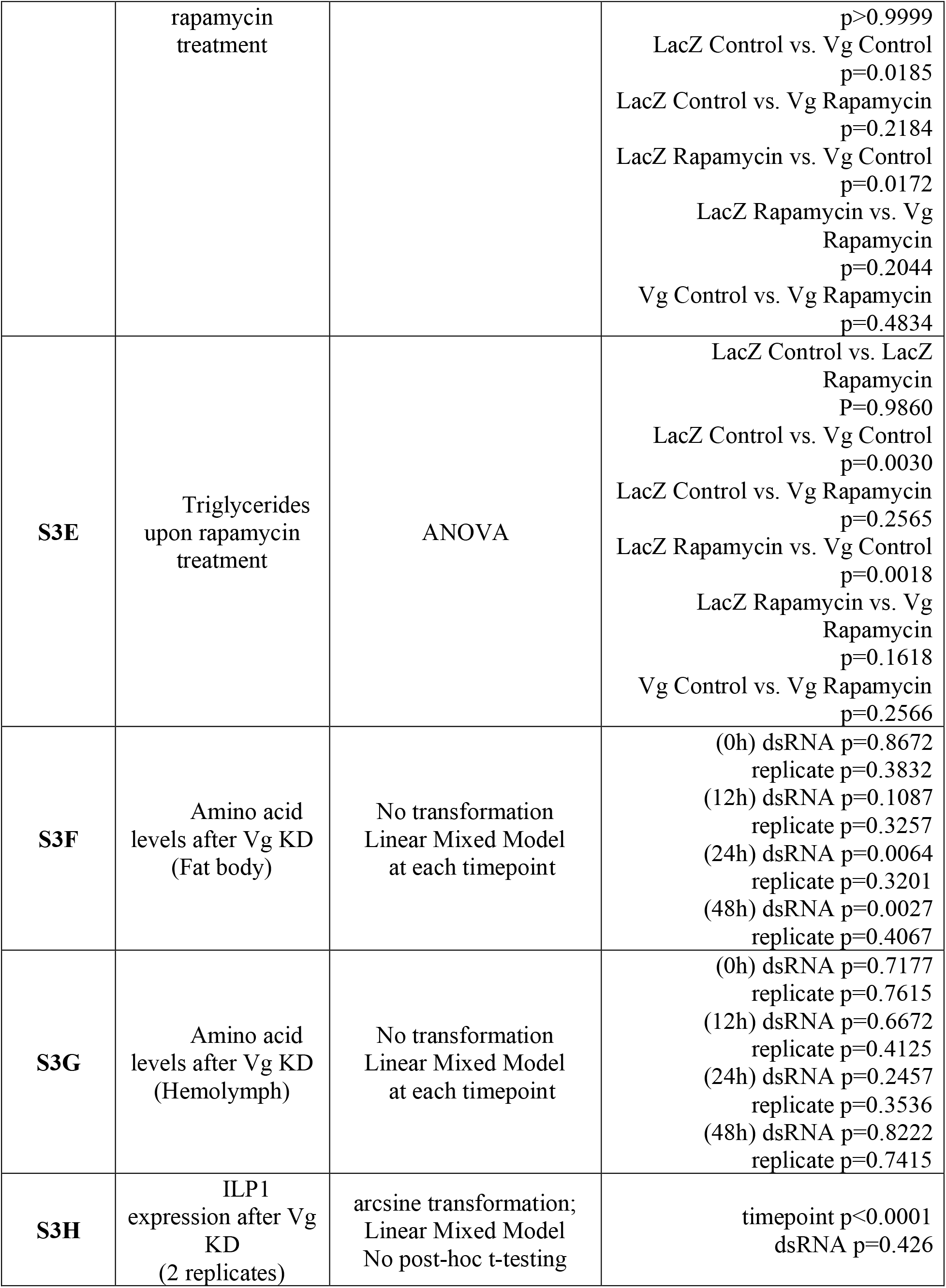
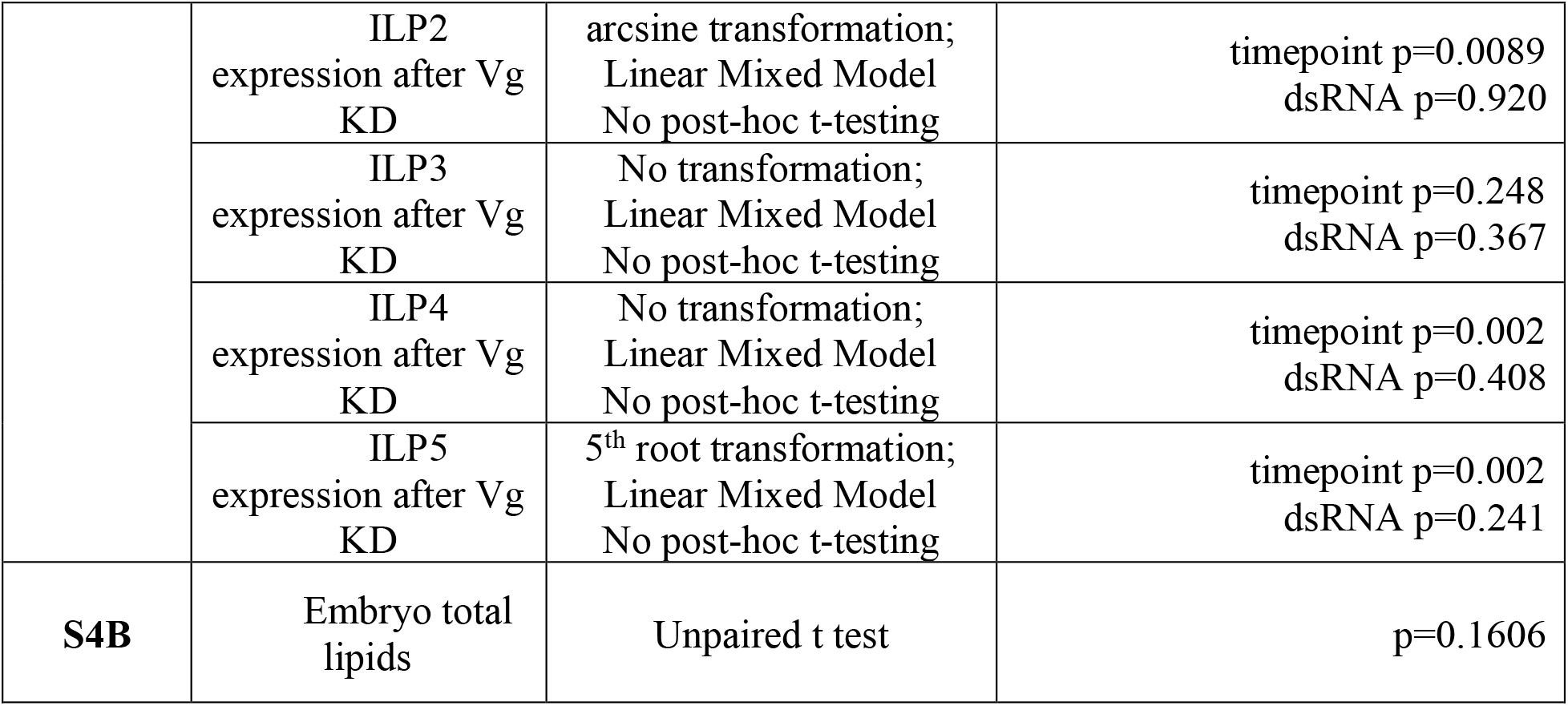
Details of statistical tests and outputs are summarized for each figure. For qRT-PCR, at least three independent biological replicates of a gene expression timecourse were analyzed, except for ILP1, where one replicate was excluded as an outlier. Effect test outputs are reported here. Multiple comparisons were calculated using pairwise Student’s t tests at each timepoint followed by FDR correction (see Table S6). KD = knock down; rand = random effect; FDR = false discovery rate.

**Supplemental Table 6.** Post-hoc testing for significant differences using an FDR of 0.05. See **Supplemental Table 5.**

| **2F**<br>**Lp after Vg KD** | **p-value** | **FDR-adjusted**<br>**p-value** | **Significant?** |
| --- | --- | --- | --- |
| dsLacZ – dsVg at 0h | 0.6414 | 0.6414 | No |
| dsLacZ – dsVg at 12h | 0.0731 | 0.0914 | No |
| dsLacZ – dsVg at 24h | $4.13 \times 10^{-6}$ | $2.07 \times 10^{-5}$ | Yes |
| dsLacZ – dsVg at 36h | 0.0020 | 0.0050 | Yes |
| dsLacZ – dsVg at 48h | 0.0450 | 0.0750 | No |
| **S1B**<br>**Vg after Lp KD** | **p-value** | **FDR-adjusted**<br>**p-value** | **Significant?** |
| dsLacZ – dsLp at 0h | 0.9198 | 0.9198 | No |
| dsLacZ – dsLp at 12h | 0.7704 | 0.9198 | No |
| dsLacZ – dsLp at 24h | 0.0029 | 0.0048 | Yes |
| dsLacZ – dsLp at 36h | 0.0028 | 0.0048 | Yes |
| dsLacZ – dsLp at 48h | $7.55 \times 10^{-5}$ | $3.78 \times 10^{-4}$ | Yes |
| **S1A**<br>**Lp after Lp KD** | **p-value** | **FDR-adjusted**<br>**p-value** | **Significant?** |
| dsLacZ – dsLp at 0h | $2.64 \times 10^{-5}$ | $6.60 \times 10^{-5}$ | Yes |
| dsLacZ – dsLp at 12h | $4.36 \times 10^{-7}$ | $2.18 \times 10^{-6}$ | Yes |
| dsLacZ – dsLp at 24h | 0.0138 | 0.0173 | Yes |
| dsLacZ – dsLp at 36h | 0.8884 | 0.8884 | No |
| dsLacZ – dsLp at 48h | $2.67 \times 10^{-4}$ | $4.45 \times 10^{-4}$ | Yes |
| **S1C (Ovaries)** | **p-value** | **FDR-adjusted p-**<br>**value** | **Significant?** |
| dsLacZ – dsVg at 0h | 0.7766 | 0.7766 | No |
| dsLacZ – dsVg at 24h | 0.6119 | 0.7766 | No |
| dsLacZ – dsVg at 48h | 0.0009 | 0.0027 | Yes |
| **S1C (Fat body)** | **p-value** | **FDR-adjusted p-**<br>**value** | **Significant?** |
| dsLacZ – dsVg at 0h | 0.7447 | 0.7447 | No |
| dsLacZ – dsVg at 24h | 0.1780 | 0.2670 | No |
| dsLacZ – dsVg at 48h | $2.86 \times 10^{-10}$ | $8.56 \times 10^{-10}$ | Yes |
| **S2A**<br>**Vg after Vg KD** | **p-value** | **FDR-adjusted**<br>**p-value** | **Significant?** |
| dsLacZ – dsVg at 0h | 0.9000 | 0.9000 | No |
| dsLacZ – dsVg at 12h | 0.0057 | 0.0115 | Yes |
| dsLacZ – dsVg at 24h | $2.94 \times 10^{-5}$ | $1.47 \times 10^{-4}$ | Yes |
| dsLacZ – dsVg at 36h | 0.0069 | 0.0115 | Yes |
| dsLacZ – dsVg at 48h | 0.5501 | 0.6877 | No |
| **S2D**<br>**Lp after Vg KD#2** | **p-value** | **FDR-adjusted**<br>**p-value** | **Significant?** |
| dsLacZ – dsVg at 0h | 0.3785 | 0.5678 | No |
| dsLacZ – dsVg at 24h | 0.0098 | 0.0294 | Yes |
| dsLacZ – dsVg at 48h | 0.8191 | 0.8191 | No |
| **S2E (Ovaries)** | **p-value** | **FDR-adjusted p-**<br>**value** | **Significant?** |
| dsLacZ – dsVg at 0h | 0.9392 | 0.9392 | No |
| dsLacZ – dsVg at 24h | 0.0210 | 0.0315 | Yes |
| dsLacZ – dsVg at 48h | 0.0177 | 0.0315 | No |
| **S2E (Fat body)** | **p-value** | **FDR-adjusted p-**<br>**value** | **Significant?** |
| dsLacZ – dsVg at 0h | 0.1927 | 0.1927 | No |
| dsLacZ – dsVg at 24h | 0.0133 | 0.0399 | Yes |
| dsLacZ – dsVg at 48h | 0.0298 | 0.0447 | Yes |
| **S3A (Fat body)** | **p-value** | **FDR-adjusted p-**<br>**value** | **Significant?** |
| dsLacZ – dsVg at 0h | 0.9058 | 0.9058 | No |
| dsLacZ – dsVg at 12h | 0.1829 | 0.3658 | No |
| dsLacZ – dsVg at 24h | $8.06 \times 10^{-25}$ | $3.22 \times 10^{-24}$ | Yes |
| dsLacZ – dsVg at 48h | 0.6257 | 0.8343 | No |

## Acknowledgements

We thank Kate Thornburg, Emily Selland, Elizabeth Nelson, Kaileigh Bumpus, and Aaron Stanton for rearing mosquitoes used in this study. We thank all other members of the Catteruccia lab for their ideas and feedback, as well as Krystle Kalafut for her insights on TOR signalling. We thank Christoph Thiele for providing the LD540 stain. Electron Microscopy Imaging, consultation and services were performed in the HMS Electron Microscopy Facility, and we thank Maria Ericsson, Peg Coughlin, and Anja Nordstrom for their help.

## Funding

F.C. is funded by the Howard Hughes Medical Institute (HHMI) as an HHMI investigator (www.hhmi.org), and by the National Institutes of Health (NIH) (R01AI148646, R01AI153404, www.nih. gov). I.S. is funded by Natural Sciences and Engineering Research Council of Canada (NSERC) as a postgraduate scholarship recipient (PGSD3 - 545866 - 2020). MI is funded by Charles A. King Trust postdoctoral research fellowship in basic science from Health Resources in Action (HRiA). The findings and conclusions within this publication are those of the authors and do not necessarily reflect positions or policies of the HHMI or the NIH. The funders had no role in the study design, in data collection, analysis or interpretation, in the decision to publish, or the preparation of the manuscript.

## Declaration of interests

The authors do not have any conflicts of interest.

## Notes

### Competing Interest Statement

The authors have declared no competing interest.

## References

Ahmed, Ashraf M, and Hilary Hurd. 2006. “Immune Stimulation and Malaria Infection Impose Reproductive Costs in Anopheles Gambiae via Follicular Apoptosis.” Microbes and Infection 8 (2): 308–15. https://doi.org/10.1016/j.micinf.2005.06.026.

Arsic, Dany, and Patrick M Guerin. 2008. “Nutrient Content of Diet Affects the Signaling Activity of the Insulin/Target of Rapamycin/P70 S6 Kinase Pathway in the African Malaria Mosquito Anopheles Gambiae.” Journal of Insect Physiology.

Atella, Georgia C, Mário Alberto C Silva-Neto, Daniel M Golodne, Shamsul Arefin, and Mohammed Shahabuddin. 2006. “Anopheles Gambiae Lipophorin: Characterization and Role in Lipid Transport to Developing Oocyte.” Insect Biochemistry and Molecular Biology 36 (5): 375–86. https://doi.org/10.1016/j.ibmb.2006.01.019.

Attardo, Geoffrey M, Immo A Hansen, and Alexander S Raikhel. 2005. “Nutritional Regulation of Vitellogenesis in Mosquitoes: Implications for Anautogeny.” Insect Biochemistry and Molecular Biology 35 (7): 661–75. https://doi.org/10.1016/j.ibmb.2005.02.013.

Blandin, Stephanie, Shin-Hong Shiao, Luis F Moita, Chris J Janse, Andrew P Waters, Fotis C Kafatos, and Elena A Levashina. 2004. “Complement-like Protein TEP1 Is a Determinant of Vectorial Capacity in the Malaria Vector Anopheles Gambiae.” Cell 116 (5): 661–70. https://doi.org/10.1016/s0092-8674(04)00173-4.

Bownes, Mary, Katrin Lineruth, and Debbie Mauchline. 1991. “Egg Production and Fertility in Drosophila Depend upon the Number of Yolk-Protein Gene Copies.” Molecular and General Genetics MGG 228 (1): 324–27. https://doi.org/10.1007/BF00282485.

Briegel, H. 1990. “Metabolic Relationship between Female Body Size, Reserves, and Fecundity of Aedes Aegypti.” Journal of Insect Physiology.

Brown, M R, R Graf, K M Swiderek, D Fendley, T H Stracker, D E Champagne, and A O Lea. 1998. “Identification of a Steroidogenic Neurohormone in Female Mosquitoes.” The Journal of Biological Chemistry 273 (7): 3967–71. https://doi.org/10.1074/jbc.273.7.3967.

Brown, Mark R, Kevin D Clark, Monika Gulia, Zhangwu Zhao, Stephen F Garczynski, Joe W Crim, Richard J Suderman, and Michael R Strand. 2008. “An Insulin-like Peptide Regulates Egg Maturation and Metabolism in the Mosquito Aedes Aegypti.” Proceedings of the National Academy of Sciences of the United States of America 105 (15): 5716–21. https://doi.org/10.1073/pnas.0800478105.

Ciudad, Laura, Maria-Dolors Piulachs, and Xavier Bellés. 2006. “Systemic RNAi of the Cockroach Vitellogenin Receptor Results in a Phenotype Similar to That of the Drosophila Yolkless Mutant.” The FEBS Journal 273 (2): 325–35. https://doi.org/https://doi.org/10.1111/j.1742-4658.2005.05066.x.

Clements, Alan. 1992. Biology of Mosquitoes, Volume 1 Development, Nutritionnd Reproduction.

Coelho, Roberta R, José Dijair Antonino de Souza Júnior, Alexandre A P Firmino, Leonardo L P de Macedo, Fernando C A Fonseca, Walter R Terra, Gilbert Engler, Janice de Almeida Engler, Maria Cristina M da Silva, and Maria Fatima Grossi-de-Sa. 2016. “Vitellogenin Knockdown Strongly Affects Cotton Boll Weevil Egg Viability but Not the Number of Eggs Laid by Females.” Meta Gene 9 (September): 173–80. https://doi.org/10.1016/j.mgene.2016.06.005.

DiMario, P J, and A P Mahowald. 1987. “Female Sterile (1) Yolkless: A Recessive Female Sterile Mutation in Drosophila Melanogaster with Depressed Numbers of Coated Pits and Coated Vesicles within the Developing Oocytes.” Journal of Cell Biology 105 (1): 199–206. https://doi.org/10.1083/jcb.105.1.199.

Dong, J, H Qiu, M Garcia-Barrio, J Anderson, and A G Hinnebusch. 2000. “Uncharged TRNA Activates GCN2 by Displacing the Protein Kinase Moiety from a Bipartite TRNA-Binding Domain.” Molecular Cell 6 (2): 269–79. https://doi.org/10.1016/s1097-2765(00)00028-9.

Feng, Yuebiao, Lu Chen, Li Gao, Li Dong, Han Wen, Xiumei Song, Fang Luo, Gong Cheng, and Jingwen Wang. 2021. “Rapamycin Inhibits Pathogen Transmission in Mosquitoes by Promoting Immune Activation.” PLoS Pathogens 17 (2): e1009353. https://doi.org/10.1371/journal.ppat.1009353.

Ford, P S, and M C Van Heusden. 1994. “Triglyceride-Rich Lipophorin in Aedes Aegypti (Diptera: Culicidae).” Journal of Medical Entomology 31 (3): 435–41. https://doi.org/10.1093/jmedent/31.3.435.

Goltsev, Yury, William Hsiong, Gregory Lanzaro, and Mike Levine. 2004. “Different Combinations of Gap Repressors for Common Stripes in Anopheles and Drosophila Embryos.” Developmental Biology 275 (2): 435–46. https://doi.org/10.1016/j.ydbio.2004.08.021.

González-Santoyo, Isaac, and Alex Córdoba-Aguilar. 2012. “Phenoloxidase: A Key Component of the Insect Immune System.” Entomologia Experimentalis et Applicata 142 (1): 1–16. https://doi.org/https://doi.org/10.1111/j.1570-7458.2011.01187.x.

Gutzeit, Herwig O, and Detlev Arendt. 1994. “Blocked Endocytotic Uptake by the Oocyte Causes Accumulation of Vitellogenins in the Haemolymph of the Female-Sterile Mutants QuitPX61 and Stand StillPS34 of Drosophila.” Cell and Tissue Research 275 (2): 291–98. https://doi.org/10.1007/BF00319427.

Hansen, Immo A, Geoffrey M Attardo, Jong-Hwa Park, Quan Peng, and Alexander S Raikhel. 2004. “Target of Rapamycin-Mediated Amino Acid Signaling in Mosquito Anautogeny.” Proceedings of the National Academy of Sciences 101 (29): 10626–31. https://doi.org/10.1073/pnas.0403460101.

Hansen, Immo A, Geoffrey M Attardo, Saurabh G Roy, and Alexander S Raikhel. 2005. “Target of Rapamycin-Dependent Activation of S6 Kinase Is a Central Step in the Transduction of Nutritional Signals during Egg Development in a Mosquito.” The Journal of Biological Chemistry 280 (21): 20565–72. https://doi.org/10.1074/jbc.M500712200.

Hansen, Immo, Geoffrey Attardo, Stacy Rodriguez, and Lisa Drake. 2014. “Four-Way Regulation of Mosquito Yolk Protein Precursor Genes by Juvenile Hormone-, Ecdysone-, Nutrient-, and Insulin-like Peptide Signaling Pathways .” Frontiers in Physiology . https://www.frontiersin.org/articles/10.3389/fphys.2014.00103.

Juhn, Jennifer, and Anthony A James. 2012. “Hybridization in Situ of Salivary Glands, Ovaries, and Embryos of Vector Mosquitoes.” Journal of Visualized Experiments : JoVE, no. 64 (June). https://doi.org/10.3791/3709.

Kim, S R, R Yao, Q Han, B M Christensen, and J Li. 2005. “Identification and Molecular Characterization of a Prophenoloxidase Involved in Aedes Aegypti Chorion Melanization.” Insect Molecular Biology 14 (2): 185–94. https://doi.org/10.1111/j.1365-2583.2004.00547.x.

Kunkel, Joseph G, and John H Nordin. 1985. “Yolk Proteins.” *Comprehensive Insect Physiology*, Biochemistry and Pharmacology, 83–111.

Li, Hongyan, and Shicui Zhang. 2017. “Functions of Vitellogenin in Eggs.” In, 389–401. https://doi.org/10.1007/978-3-319-60855-6_17.

Lin, Ying, Yan Meng, Yan-Xia Wang, Juan Luo, Susumu Katsuma, Cong-Wen Yang, Yutaka Banno, Takahiro Kusakabe, Toru Shimada, and Qing-You Xia. 2013. “Vitellogenin Receptor Mutation Leads to the Oogenesis Mutant Phenotype ‘Scanty Vitellin’ of the Silkworm, Bombyx Mori.” The Journal of Biological Chemistry 288 (19): 13345–55. https://doi.org/10.1074/jbc.M113.462556.

Marcenac, Perrine, W Robert Shaw, Evdoxia G Kakani, Sara N Mitchell, Adam South, Kristine Werling, Eryney Marrogi, et al. 2020. “A Mating-Induced Reproductive Gene Promotes Anopheles Tolerance to Plasmodium Falciparum Infection.” PLoS Pathogens 16 (12): e1008908. https://doi.org/10.1371/journal.ppat.1008908.

Marinotti, Osvaldo, Margareth de L Capurro, Xavier Nirmala, Eric Calvo, and Anthony A James. 2006. “Structure and Expression of the Lipophorin-Encoding Gene of the Malaria Vector, Anopheles Gambiae.” *Comparative Biochemistry and Physiology. Part B*, Biochemistry & Molecular Biology 144 (1): 101–9. https://doi.org/10.1016/j.cbpb.2006.01.012.

Miraldi, Emily R, Hadar Sharfi, Randall H Friedline, Hannah Johnson, Tejia Zhang, Ken S Lau, Hwi Jin Ko, et al. 2013. “Molecular Network Analysis of Phosphotyrosine and Lipid Metabolism in Hepatic PTP1b Deletion Mice.” Integrative Biology : Quantitative Biosciences from Nano to Macro 5 (7): 940–63. https://doi.org/10.1039/c3ib40013a.

Mitchell III, Robert D, Elizabeth Ross, Christopher Osgood, Daniel E Sonenshine, Kevin V Donohue, Sayed M Khalil, Deborah M Thompson, and R Michael Roe. 2007. “Molecular Characterization, Tissue-Specific Expression and RNAi Knockdown of the First Vitellogenin Receptor from a Tick.” Insect Biochemistry and Molecular Biology 37 (4): 375–88. https://doi.org/https://doi.org/10.1016/j.ibmb.2007.01.005.

Park, Jong-Hwa, Geoffrey M Attardo, Immo A Hansen, and Alexander S Raikhel. 2006. “GATA Factor Translation Is the Final Downstream Step in the Amino Acid/Target-of-Rapamycin-Mediated Vitellogenin Gene Expression in the Anautogenous Mosquito Aedes Aegypti.” The Journal of Biological Chemistry 281 (16): 11167–76. https://doi.org/10.1074/jbc.M601517200.

Peng, Lu, Qing Wang, Ming-Min Zou, Yu-Dong Qin, Liette Vasseur, Li-Na Chu, Yi-Long Zhai, et al. 2020. “CRISPR/Cas9-Mediated Vitellogenin Receptor Knockout Leads to Functional Deficiency in the Reproductive Development of Plutella Xylostella .” Frontiers in Physiology . https://www.frontiersin.org/articles/10.3389/fphys.2019.01585.

Qin, Pu, Pelin Arabacilar, Roberta E Bernard, Weike Bao, Alan R Olzinski, Yuanjun Guo, Hind Lal, et al. 2017. “Activation of the Amino Acid Response Pathway Blunts the Effects of Cardiac Stress.” Journal of the American Heart Association 6 (5). https://doi.org/10.1161/JAHA.116.004453.

Raikhel, A S, and T S Dhadialla. 1992. “Accumulation of Yolk Proteins in Insect Oocytes.” Annual Review of Entomology 37: 217–51. https://doi.org/10.1146/annurev.en.37.010192.001245.

Rogers, David W, Miranda M A Whitten, Janis Thailayil, Julien Soichot, Elena A Levashina, and Flaminia Catteruccia. 2008. “Molecular and Cellular Components of the Mating Machinery in Anopheles Gambiae Females.” Proceedings of the National Academy of Sciences 105 (49): 19390–95. https://doi.org/10.1073/pnas.0809723105.

Rono, Martin K., Miranda M. A. Whitten, Mustapha Oulad-Abdelghani, Elena A. Levashina, and Eric Marois. 2010. “The Major Yolk Protein Vitellogenin Interferes with the Anti-Plasmodium Response in the Malaria Mosquito Anopheles Gambiae.” Edited by David S. Schneider. PLoS Biology 8 (7): e1000434. https://doi.org/10.1371/journal.pbio.1000434.

ROTH, T F, and K R Porter. 1964. “YOLK PROTEIN UPTAKE IN THE OOCYTE OF THE MOSQUITO AEDES AEGYPTI. L.” The Journal of Cell Biology 20 (2): 313–32. https://doi.org/10.1083/jcb.20.2.313.

Shang, F, J-Z Niu, B-Y Ding, Q Zhang, C Ye, W Zhang, G Smagghe, and J-J Wang. 2018. “Vitellogenin and Its Receptor Play Essential Roles in the Development and Reproduction of the Brown Citrus Aphid, Aphis (Toxoptera) Citricidus.” Insect Molecular Biology 27 (2): 221–33. https://doi.org/10.1111/imb.12366.

Shaw, W Robert, Eleonora Teodori, Sara N Mitchell, Francesco Baldini, Paolo Gabrieli, David W Rogers, and Flaminia Catteruccia. 2014. “Mating Activates the Heme Peroxidase HPX15 in the Sperm Storage Organ to Ensure Fertility in Anopheles Gambiae.” Proceedings of the National Academy of Sciences 111 (16): 5854–59. https://doi.org/10.1073/pnas.1401715111.

Sikalidis, Angelos K. 2013. “Cellular and Animal Indispensable Amino Acid Limitation Responses and Health Promotion. Can the Two Be Linked? A Critical Review.” International Journal of Food Sciences and Nutrition 64 (3): 300–311. https://doi.org/10.3109/09637486.2012.738649.

Spandl, Johanna, Daniel J White, Jan Peychl, and Christoph Thiele. 2009. “Live Cell Multicolor Imaging of Lipid Droplets with a New Dye, LD540.” *Traffic (Copenhagen*, Denmark) 10 (11): 1579–84. https://doi.org/10.1111/j.1600-0854.2009.00980.x.

Sun, Jianxin, Tsuyoshi Hiraoka, Neal T Dittmer, Kook-Ho Cho, and Alexander S Raikhel. 2000. “Lipophorin as a Yolk Protein Precursor in the Mosquito, Aedes Aegypti.” Insect Biochemistry and Molecular Biology 30 (12): 1161–71. https://doi.org/https://doi.org/10.1016/S0965-1748(00)00093-X.

Valvezan, Alexander J, and Brendan D Manning. 2019. “Molecular Logic of MTORC1 Signalling as a Metabolic Rheostat.” Nature Metabolism 1 (3): 321–33. https://doi.org/10.1038/s42255-019-0038-7.

Vlachou, Dina, Timm Schlegelmilch, George K Christophides, and Fotis C Kafatos. 2005. “Functional Genomic Analysis of Midgut Epithelial Responses in Anopheles during Plasmodium Invasion.” Current Biology : CB 15 (13): 1185–95. https://doi.org/10.1016/j.cub.2005.06.044.

Wang, Jing, Shasha Yu, Luhan Wang, Tingting Liu, Xuesen Yang, Xiaobing Hu, and Ying Wang. 2022. “Capsaicin Decreases Fecundity in the Asian Malaria Vector Anopheles Stephensi by Inhibiting the Target of Rapamycin Signaling Pathway.” Parasites & Vectors 15 (1): 458. https://doi.org/10.1186/s13071-022-05593-0.

Werling, Kristine, W. Robert Shaw, Maurice A. Itoe, Kathleen A. Westervelt, Perrine Marcenac, Douglas G. Paton, Duo Peng, et al. 2019. “Steroid Hormone Function Controls Non-Competitive Plasmodium Development in Anopheles.” Cell 177 (2): 315–325.e14. https://doi.org/10.1016/j.cell.2019.02.036. World Malaria Report 2022. 2022.

